# Maximizing meiotic crossover rate reveals the map of Crossover Potential

**DOI:** 10.1101/2024.11.05.622098

**Authors:** Juli Jing, Qichao Lian, Stephanie Durand, Raphael Mercier

## Abstract

Meiotic crossovers are limited in number and unevenly distributed along chromosomes, both features often differing between sexes. The mechanisms imposing a different crossover landscape in female and male meiosis remain elusive. Here, we simultaneously disrupted multiple anti-crossover mechanisms in Arabidopsis and analyzed the whole genome sequence of thousands of female- and male-derived progenies. The largest crossover increase was reached in *zyp1 recq4*, with 12-fold in females and a 4.5-fold increase in males. Despite this unprecedented level of crossovers, fertility is marginally affected, opening new possibilities for plant breeding. Manipulating additional crossover regulators in *zyp1 recq4* did not further elevate the frequency of crossovers, but modified the relative contributions of the two known crossover pathways. This suggests an upper limit was reached and the two pathways compete for a large but limited set of recombination intermediates. Remarkably, while wild-type crossover distribution differs markedly between sexes, the crossover landscapes of diverse mutants in both females and males converge to a single novel profile, which we termed Crossover Potential (CO_P_). The CO_P_ profile, which we defined using 49,482 crossovers, can be accurately predicted using only sequence divergence and chromatin features. We propose that the CO_P_ represents the density of eligible recombination precursors, which is determined by genomic features and is thus identical in females and males. It suggests that the sexual dimorphism in the crossover landscape results exclusively from differential regulation of the likeness of precursors to mature into crossovers.

## Introduction

In sexual reproduction, meiotic recombination generates reciprocal exchanges between homologous chromosomes called crossovers (CO), enhancing genetic diversity ^1,2^. Notably, in the context of plant breeding, COs introduce new favorable gene combinations and break up unfavorable linkages, allowing breeders to improve crop varieties. However, COs are limited in number, with typically 1-3 per chromosome per meiosis ^3,4^, despite a large excess of molecular precursors. CO formation is initiated by programmed DNA double-strand breaks (DSBs) which are repaired by using the homologous chromosome as a template. The number of early recombination intermediates is estimated to be 10-25 fold greater than the eventual number of COs in mammals and plants ^5–9^. CO frequencies are not homogenous along chromosomes, creating a landscape of recombination with high peaks and large valleys. COs are also subjected to interference, in which one CO inhibits the formation of another nearby, effectively limiting the frequency of close double COs ^10–12^. Intriguingly, CO frequency and distribution can differ markedly between females and males of the same species, a phenomenon called heterochiasmy ^13^. The factors and mechanisms that shape CO distribution along the genome and differently in females and males are not well understood. In particular, it is unclear what the relative contribution of precursor distribution versus a differential likelihood of precursor maturation to CO is in the final landscape.

In most eukaryotes, two molecular pathways contribute to the maturation of recombination intermediates into CO, defining two classes of COs. Class I COs are promoted by the ZMMs protein, are sensitive to interference, and account for the majority of COs; class II COs are ZMM-independent and promoted by nucleases including MUS81. Class II is a minor pathway in most eukaryotes including plants, and is not or much less sensitive to interference ^11,14^.

A series of factors that limit either class I or class II COs are known in Arabidopsis. One is the dosage of the ZMM protein HEI10, whose overexpression (HEI10^oe^) almost doubles the number of class I COs ^15,16^. Another key factor regulating class I CO is the central part of the synaptonemal complex (SC), including the transverse filament ZYP1 ^17,18^ ^19^. The SC forms a structure that connects the homologous chromosomes all their length during the meiotic prophase. Mutation of ZYP1 eliminates CO interference and heterochiasmy and increases class I CO numbers ∼twice ^17,18^ and has a cumulative effect with HEI10^oe^ on the number of class I CO ^20^. On the other hand, the RECQ4A and RECQ4B helicases (called here RECQ4 for simplicity) redundantly prevent the formation of class II CO ^3,21^. The combination of *recq4* mutation and HEI10^oe^ showed an additive effect with an increase in both class I and class II COs ^15^, suggesting that class I and class II CO can be manipulated independently. The AAA-ATPase protein FIGL1 also limits class II COs by regulating strand invasion ^22,23^ with has a synergetic effect with *recq4*, reaching the largest increase of (class II) CO described to date ^3,22,24,25^. Additional proteins, such as FLIP, SNI, TOP3/RMI1, HSBP, HCR1 and HCR3 have anti-CO roles in Arabidopsis, being partners or regulators of the functions described above ^24,26–30^. Several combinations of anti-CO factors including higher-order ones, remained to be tested to explore the upper limit of crossover frequencies.

## Results

### HEI10^oe^*, zyp1* and *recq4* act differently in boosting CO numbers

We analyzed the number and distribution of COs in both female and male meiosis in wild-type, single and multi-mutants, by generating biallelic-Col/L*er* F1 hybrids and reciprocally crossing them with wild-type Col (**Fig. 1a, Fig. S1-S4**). The obtained female and male-derived progenies were analyzed by whole genome Illumina sequencing to detect CO transmitted by female and male gametes, respectively (**Fig. 1b-c**). In parallel, we counted the number of MLH1 foci, which marks class I COs, in female and male meiocytes of Col/L*er* F1 hybrids of each genotype (**Fig. 2a-m**). Comparison of the number of MLH1 foci (class I CO) and genetic crossovers detected by sequencing (class I + class II COs), allows the estimation of the contribution of each pathway (**Fig. 2n**). Note that the mean number of CO per gamete is expected to be half the number of cytological CO (or chiasma). This is because a chiasma affects only two of the four chromatids of a chromosome pair, and a gamete inherits a single chromatid ^31^ (see figure S11 in ^20^). In **Fig. 2n**, the number of genetic CO expected to be observed if each MLH1 foci is converted into a CO is indicated by a dashed line (1 foci=0.5 CO).

**Figure 1:**
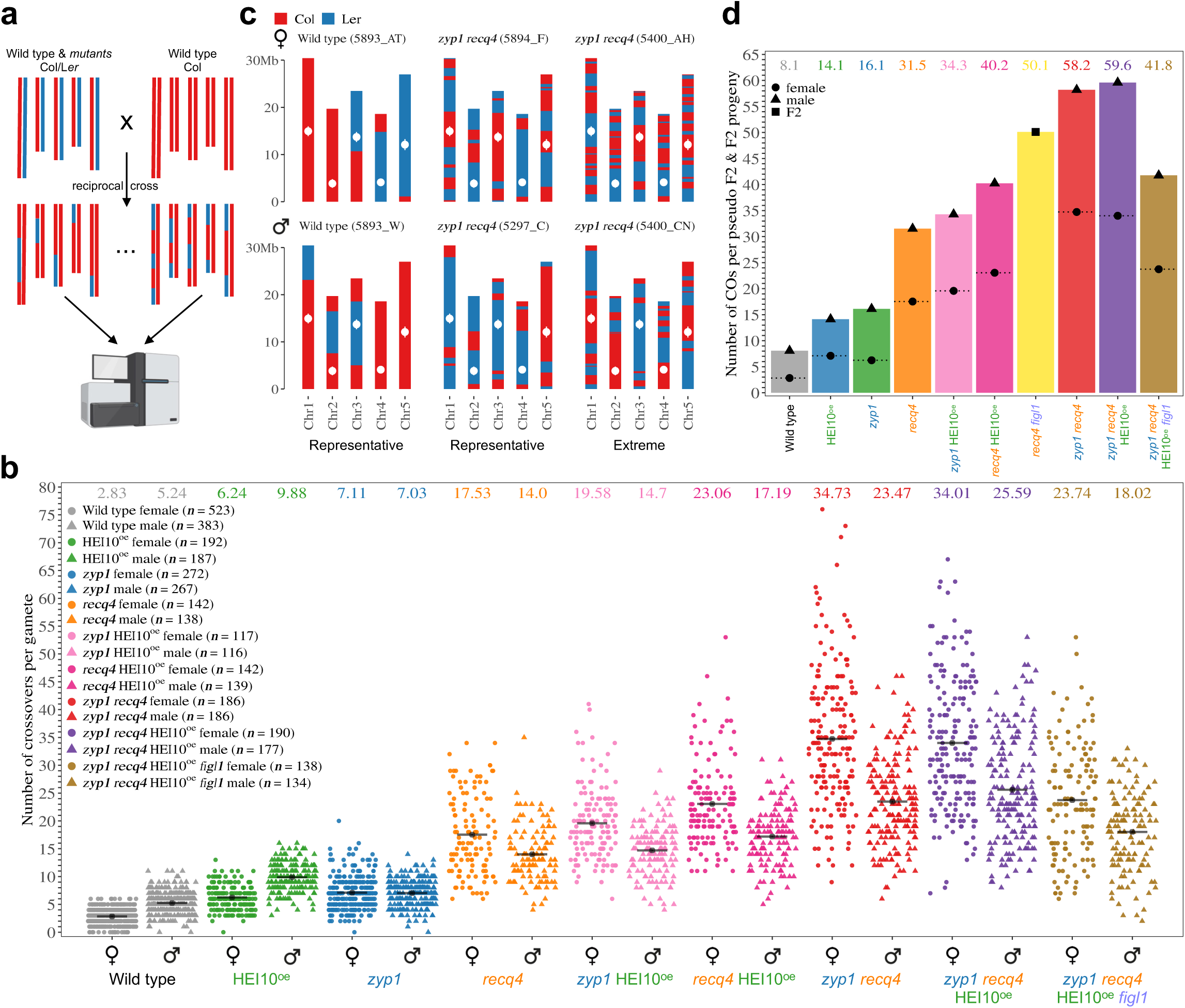
Combination of *zyp1* and *recq4* massively increases meiotic crossover frequency. **a.** Schematic representation of the experimental design. Col/L*er* hybrid plants, either wild-type or carrying mutations, were crossed with wild-type Col in both directions. The resulting plant populations (BC1) were sequenced to score the COs transmitted by the female and male gametes of the hybrids. **b.** The number of COs per female and male gametes for each genotype. Each point represents the number of COs in one BC1/gamete. Circles and triangles are females and males, respectively. A black line indicates the mean and its value is specified at the top of the graph. The BC1/gamete population size (n) is indicated for each genotype. **c.** Representative transmitted chromatid sets are shown for selected samples. The genotype and the name of the sample are specified. Blue and red indicate Col and L*er* genotypes, a transition marking a crossover. White points indicate centromeres. **d.** The average number of COs per F2 sample (for *recq4 figl1,* yellow) or pseudo-F2 (sum of female and male averages, for all other genotypes).

**Figure 2.**
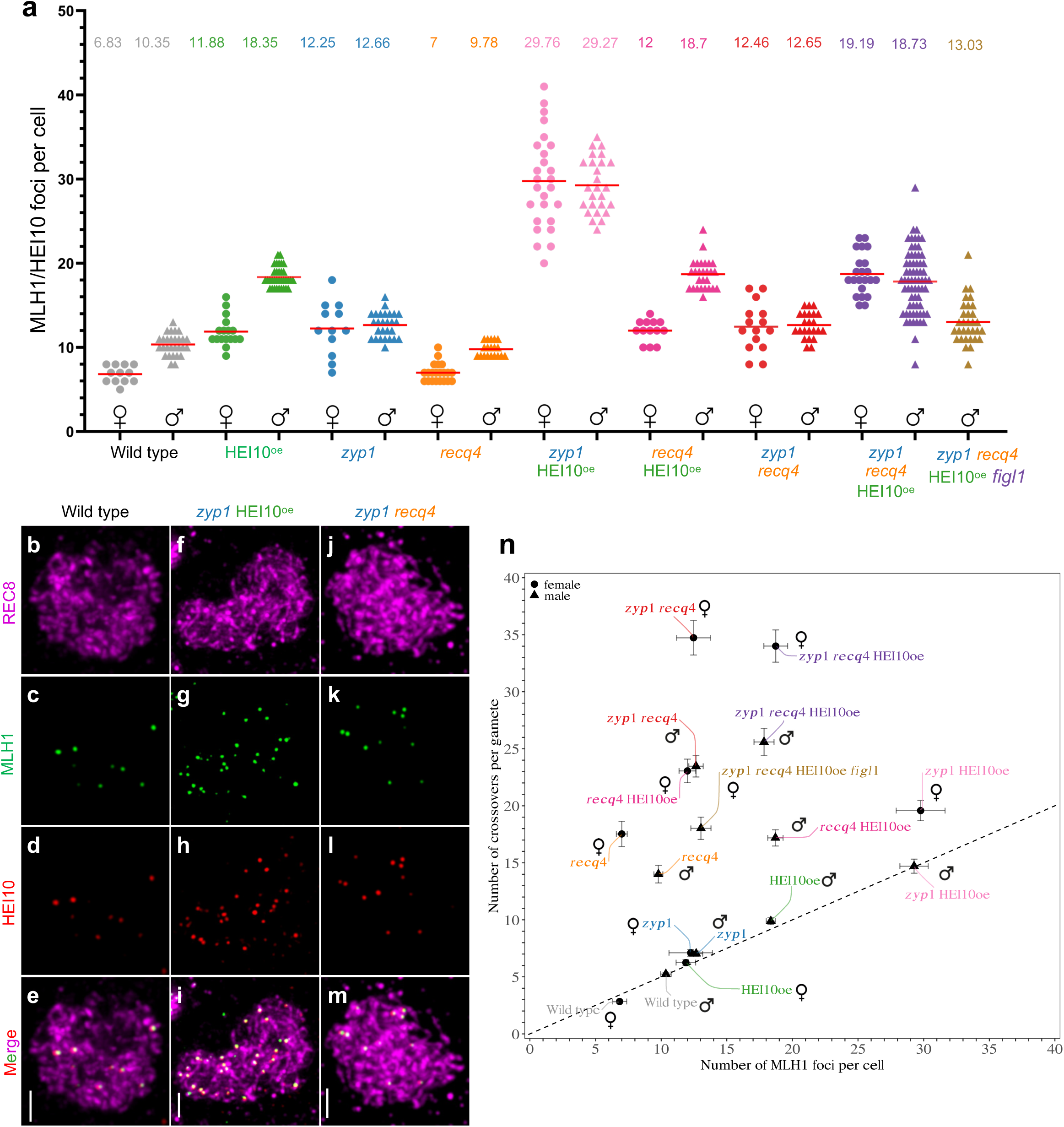
Analysis of MLH1/HEI10 foci and contributions of crossover pathways. **a.** MLH1/HEI10 foci count in male and female meiocytes of wild type and mutants. Each dot is an individual cell. Red bars and numbers indicate the mean number of MLH1/HEI10 co-foci. Circles and triangles are females and males, respectively. **b-m.** Immunostaining of REC8 (purple), MLH1 (green) and HEI10 (red) in wild type (b-e), *zyp1* HEI10^oe^ (f-i) and *zyp1 recq4*(j-m) male diplotene meiocytes. MLH1-HEI10 co-foci indicate class I CO sites. Scale bar=2 µm. **n.** Comparison of the numbers of genetic COs (Fig1a) and MLH1/HEI10 foci per cell (fig. 2a) in females and males of various genotypes. Females are shown by circle, and males are shown by triangle. Error bars are the 90% confidence intervals of the mean. The dash line represents COs per gamete is on average one-half of COs per meiocyte.

In the wild type, we observed an average of 2.8 CO per female gamete, and 5.2 per male, confirming the previously observed heterochiasmy ^32,33^. The number of MLH1 foci matches 2-fold the number of genetic CO, suggesting that in both female and male wild-type, the vast majority of crossovers are class I CO, with a negligible number of class II CO.

In single mutants, HEI10^oe^ and *zyp1*, the average CO number per transmitted gamete was substantially increased compared with the wild type, consistent with previous results ^16,17^. In HEI10^oe^, the CO number in females (6.2, 2.2-fold/wild type) remains lower than in males (9.9, 1.9-fold/wild type). In *zyp1*, the CO number is equal in female (7.1, 2.5-fold/wild type) and male meiosis (7.0, 1.3-fold/wild type). In both cases, MLH1 foci were also increased, matching the number of CO (**Fig. 2a, 2n**). Thus, overexpressing HEI10 or mutating *ZYP1* exclusively increases class I CO. However, while heterochiasmy and CO interference are still present in HEI10^oe^, both are abolished in *zyp1* (**Fig. 1b, Fig. S5**).

Single *recq4* mutation increased CO to higher levels and, remarkably, inverted heterochiasmy, reaching 17.5 in females (6.3-fold/wild type) and 14 in male gametes (4.5-fold/wild type) (**Fig. 1b**) ^3^. In contrast to HEI10^oe^ and *zyp1,* the number of MLH1 foci was unchanged in both female or male meiocytes compared to wild-type (**Fig. 2**), consistent with the role of RECQ4 in limiting specifically class II COs ^21^. This is reflected in **Fig. 2n**, by a vertical deviation from the dashed line, with the number of genetic CO largely exceeding the expected number according to MLH1 foci counts. Accordingly, as class II COs are not subjected to interference, CO interference is not detectable in *recq4* (**Fig. S5**). Thus, *recq4* boosts class II CO, with a more prominent effect in female than male meiosis.

### Maximizing CO numbers

HEI10^oe^ increases class I COs while *recq4* increases class II COs. Combining *recq4* HEI10^oe^, we observed an average of 23 COs in female and 17.2 in male gametes, higher than both single mutants (**Fig. 1b**). This is consistent with previous studies performed in F2 population ^15^. Under the hypothesis of additive effects, simply summing the wild-type counts with the gains observed in each single mutant, we would have expected 20.9 COs in female gametes [2.8+(6.2-2.8) + (17.5-2.8)] and 18.7 in male gametes [5.2+(9.9-5.2) + (14.0-5.2)], which is similar to the observed values. In both female and male meiosis, the numbers of MLH1 foci in *recq4* HEI10^oe^ are not modified compared to HEI10^oe^ (**Fig. 2a, n**), suggesting that class I COs are unaffected by the *recq4* mutation. Altogether, this shows that combining *recq4* and HEI10^oe^ leads to parallel increases of class I CO and class II provoked by respectively by HEI10^oe^ and *recq4* and thus to an additive effect on the total number of COs.

Next, we combined the two factors that individually increase class I CO, HEI10^oe^ and *zyp1*^20^. Following the same logic as above, the hypothesis of an additive effect predicts 11.7 COs in female (2.8+3.4+4.3) and 10.5 COs in male gametes (5.2+4.7+1.8). In *zyp1* HEI10^oe^, we observed an average of 14.7 COs in male gametes, which is higher than the additive prediction, and 19.6 in females, which is almost twice the prediction. MLH1 foci are largely increased in female and male meiosis, reaching ∼29.5 in both sexes, more than each single mutant and 3-4-fold the wild-type (**Fig. 2a**). This indicates that HEI10^oe^ and *zyp1* synergistically augment class I CO ^20^. In male *zyp1* HEI10^oe^ meiocytes, the number of MLH1 foci (29.3) matches the frequency of genetic COs (**Fig. 1b, Fig. 2a, n**), suggesting that all COs in male *zyp1* HEI10^oe^ are of class I. In contrast, in female *zyp1* HEI10^oe^, the MLH1 foci counts were 29.8, which corresponds to 14.9 COs per gamete, while 19.6 COs were observed (**Fig. 1b, Fig. 2a, n**). This suggests that in addition to the large boost in class I COs, class II COs are also increased in *zyp1* HEI10^oe^ female meiosis, leading to an inversion of heterochiasmy.

In the last combination of two factors, *zyp1 recq4,* the COs number per gamete was massively increased to 34.7 in females and 23.5 in males, almost twice the number measured in *recq4*, the highest single mutant (**Fig. 1b**). This is significantly higher than the predictions under an additive hypothesis, 21.8 in females [2.8 + (7.1-2.8) + (17.5-2.8)] and 15.8 in males [5.2 + (7.0-5.2) + (14-5.2)], indicating a synergetic effect. The high CO levels in *zyp1 recq4* are unprecedented (compared to wild-type, 12.4-fold in females and 4.5-fold in males), being above all previously described mutants, including the so-far champion *recq4 figl1* (**Fig. 1b-d**) ^15–17,20,21,25^. The variance of CO per gamete is also large, with the extreme case of a female gamete with 76 COs distributed among the five chromosomes, while the highest number observed in a wild-type female gamete is 6 COs (**Fig. 1b, c**). In *zyp1 recq4,* the MLH1 foci numbers (12.5 in females, 12.7 in males) are stable compared with *zyp1* (12.3 in females, 12.7 in males) (**Fig.2a, n**) and do not match the number of COs, suggesting that the vast majority of CO are class II CO (**Fig. 2a, n**). Mutating *RECQ4* in the *zyp1* mutant thus increases class II CO, like it does in wild-type, but with a much stronger effect (**Fig 2n**). Altogether, we conclude that in *zyp1 recq4* class I COs are increased similarly to the single *zyp1* while class II crossovers are boosted by a synergistic effect of the two mutations, which is particularly marked in female meiosis. This also points to a role of ZYP1 in preventing class II CO, in addition to its described role in regulating class I CO.

The results described above show that all the combinations of two mutations among HEI10^oe^, *zyp1,* and *recq4* have more COs than the single mutants, predicting that the triple mutant should have even more COs. However, the CO numbers in *zyp1 recq4* HEI10^oe^ did not increase significantly compared to *zyp1 recq4*, reaching 34.0 COs in females and 25.6 in males, respectively (**Fig. 1b**), indicating that a certain upper limit might have been reached. Interestingly, the number of MLH1 foci is increased when adding HEI10^oe^ to *zyp1 recq4*, but not the total number of COs, suggesting that an increase in class I COs is compensated by a decrease in class II CO. This further supports the conclusion that an upper limit has been reached and that the two pathways compete for a large but limited number of CO precursors.

FIGL1 is another anti-class II CO factor that acts independently from RECQ4 ^3,25^. Adding *figl1* mutation to *zyp1 recq4* HEI10^oe^ is thus expected to increase COs further. However, the number of COs in *zyp1 recq4* HEI10^oe^ *figl1* mutant was not increased and instead was decreased compared to *zyp1 recq4* HEI10^oe^ (**Fig. 1b, d**). This further supports the idea of an upper limit in CO formation and shows that the *figl1* mutation can increase or decrease COs depending on the context.

### Inverted heterochiasmy in hyper-recombinant mutants

Heterochiasmy is defined as a different rate of meiotic recombination between the two sexes of the same species ^34^. In wild-type Arabidopsis, the frequency of CO is higher in males (ratio female/ male = 0.54), with a similar ratio in HEI10^oe^ (0.63). In *zyp1*, heterochiasmy is abolished with similar CO frequency in female and male gametes (**Fig. 1b**). Intriguingly, the heterochiasmy is inverted, with more CO in females than males in single *recq4* (1.25) and all double and triple mutants studied here: *zyp1* HEI10^oe^ (1.33), *recq4* HEI10^oe^ (1.34)*, zyp1 recq4* (1.48), *zyp1 recq4* HEI10^oe^ (1.33), and *zyp1 recq4* HEI10^oe^ *figl1* (1.32) (**Fig. 1b**). This shows that while CO levels are higher in males than in females in wild-type, the potential for CO formation is higher in females.

### Fertility and meiosis are only marginally affected in *zyp1 recq4*

We estimated the fertility by counting the number of seeds per fruit (silique). Different mutants showed varying degrees of reduction in fertility, but the fertility is poorly correlated with CO frequency (**Fig. 1b, Fig. 3a**). The fertility of *recq4* was not significantly reduced compared to the wild type, despite a massive increase in COs (6.3-fold in females, 2.7-fold in males), consistent with previous results ^3,15^ (Fisher’s LSD test, p>0.9999, **Fig. 1b, Fig. 3a**). In *zyp1*, fertility is modestly but significantly affected (p<0.0001, **Fig. 3a**), probably because of the failure to ensure the obligate CO ^17,18^. Strikingly, *zyp1 recq4* showed a fertility similar to *zyp1* (p=0.9913, **Fig. 3a, b**), showing that the dramatic increase in COs is not associated with a further reduction in fertility. Despite a ∼12-fold elevated CO frequency in female meiosis, the fertility of *zyp1 recq4* is still ∼75% of wild-type (**Fig. 3a, b**). Further, analyses based on genome coverage by sequencing did not detect any aneuploids in the progeny of *zyp1 recq4* (n = 2×186) suggesting that chromosome missegregation at meiosis is absent or rare (**Fig. 3c**). However, chromosome spreads of male meiocytes revealed abnormal chromosome structures in ∼16% of metaphase I cells, suggestive of unrepaired recombination intermediates, which might cause reduced fertility in *zyp1 recq4* (**Fig. 3e, f, k, l, p**).

**Figure 3.**
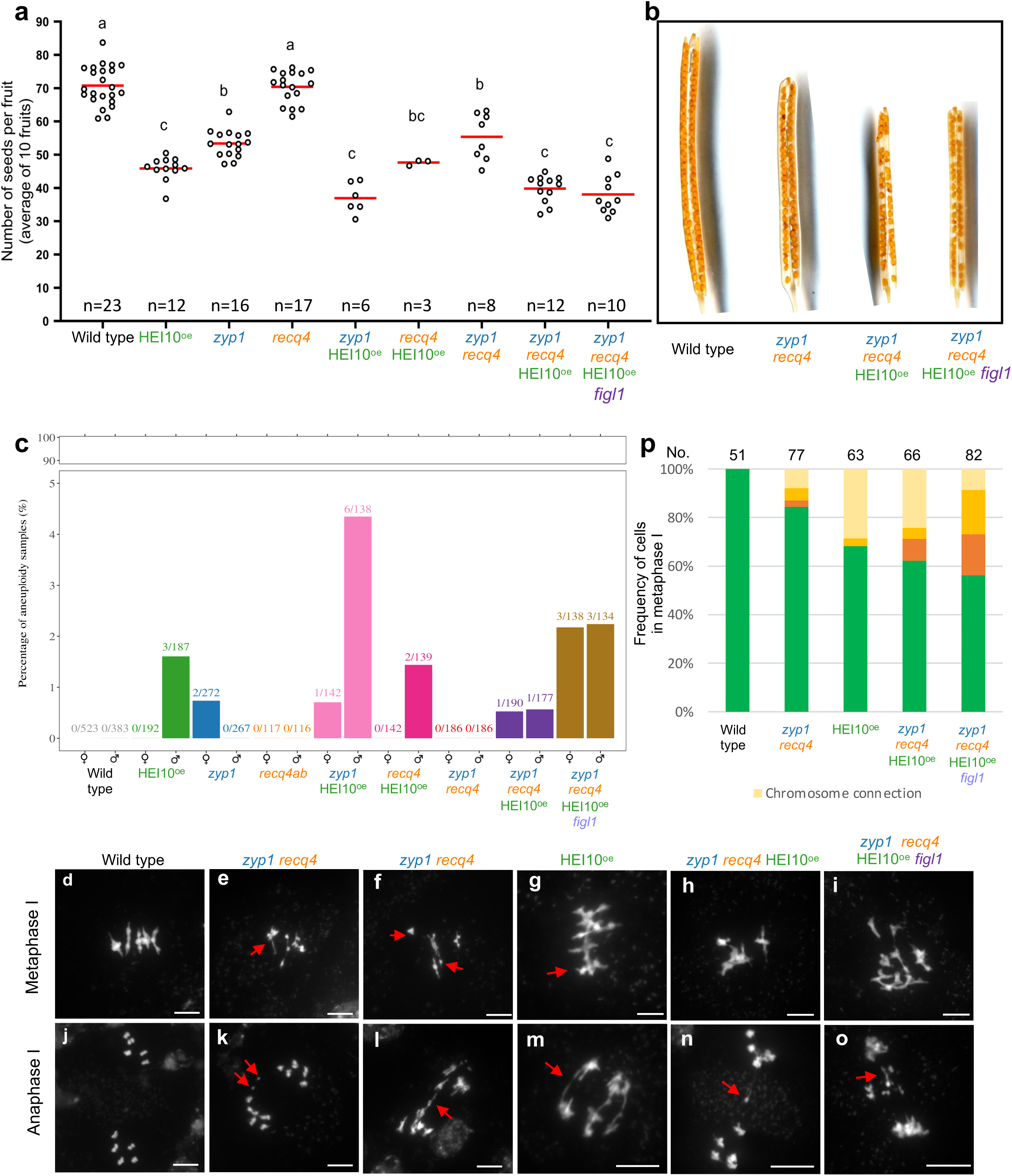
Analysis of fertility and meiosis defect. **a.** Quantification of fertility. Each dot represents the fertility of an individual plant, measured as the number of seeds per fruit averaged on ten fruits. The red bar shows the mean. All plants were grown in parallel, and the wild-type controls were siblings of the mutants. The number n of analyzed plants is indicated and p values are one-way ANOVA followed by Fisher’s LSD test. **b.** Representative cleared fruits of wild type, *zyp1 recq4*, *zyp1 recq4* HEI10^oe^ and *zyp1 recq4* HEI10^oe^ *figl1* mutants in Col/L*er* background. **c.** The percentage of aneuploid samples detected in each population. The proportion of aneuploid samples in each population is shown on top of the bars. **d-o**. DAPI-stained meiotic chromosome spreads from Col/L*er* male meiocytes in wild type (**d, j**), *zyp1 req4* (**e, f, k, i**), HEI10^oe^ (**g, m**), *zyp1 recq4* HEI10^oe^ (**h, n**) and *zyp1 recq4* HEI10^oe^ *figl1* (**i, o**). **d-i** Metaphase I. **j-o** Anaphase I. Scale bar = 10 µm. Red arrows pointed out abnormal chromosome connections, fragments and chromosome threads. **p.** Quantification of different chromosome behaviors at metaphase I in wild type, *zyp1 recq4*, HEI10^oe^, *zyp1 recq4* HEI10^oe^ and *zyp1 recq4* HEI10^oe^ *figl1*. Cells were categorized according to normal (5 bivalents) and abnormal chromosome behavior (fragmentation, univalent and chromosome connection). The number of analyzed cells is indicated above the bar.

HEI10^oe^ showed a significant fertility reduction (p<0.0001, **Fig. 3a**), consistent with previous studies ^15^, possibly due to a chromosomal rearrangement associated with the transgene in the HEI10^oe^ C2 line ^16^. Consistently chromosome connections were observed in 34% of HEI10^oe^ meiotic cells (**Fig. 3g, m, p**). Aneuploids were also detected in the progeny of all genotypes with HEI10^oe^ (**Fig. 3c**) whereas no aneuploids were detected in *recq4* and *zyp1 recq4* progenies (**Fig. 3c**). Fertility of *zyp1 recq4* HEI10^oe^ *figl1* was not reduced compared with *zyp1 recq4* HEI10^oe^ (p=0.9961, **Fig. 3a, b**), suggesting that plants can still maintain a certain level of fertility even with disturbed strand-invasion and high recombination.

### Meiotic chromosome fragmentation and reduced fertility in *zyp1 mus81*

As shown above, mutating *ZYP1* in certain contexts (*recq4*, female HEI10^oe^) provokes an increase in COs but not MLH1 foci, suggesting an anti-class II CO function for ZYP1, in addition to its established role in regulating class I COs. Mutating the nuclease MUS81 in anti-class II factors in Arabidopsis provoked meiotic catastrophe ^21,22,35,36^, presumably because DNA joint molecules remain unrepaired. Similarly, we observed some chromosome fragments in 40% of anaphase I meiotic cells in *zyp1 mus81* (**Fig. S6**). In addition, fertility in *zyp1 mus81* was reduced by 25% compared to wild type or *mus81 (***Fig. S6j).** This suggests that ZYP1 prevents the formation of joint molecules that need MUS81 to be repaired. However, this role is likely minor or redundant as no increase of class II COs is observed in the single *zyp1* mutant (**Fig. 2n**) ^17^.

### Revealing the distribution of Crossover Potential

Next, we examine the distribution of COs along the chromosomes to test whether or not the increase observed in the different mutants is homogeneous (**Fig. 4**). In all the mutant combinations with large CO increases, COs tend to accumulate in distal regions in both females and males. It is especially striking when looking at the highest recombining genotypes *zyp1 recq4* and *zyp1 recq4* HEI10^oe^ (**Fig. 4a-b**). The same is observed when merging the female and male data (pseudo-F2), and in the previously analyzed *recq4 figl1* F2 populations (**Fig. 4c**) ^25^. Intriguingly, the crossover landscapes in the different hyper-recombining mutants were highly consistent with each other, with common peaks and valleys (see also below). Looking at the local fold increase compared to the wild type (**Fig. S7**), we detected common hotspots of CO increase in mutants, some above 200-fold in *zyp1 recq4* and *zyp1 recq4* HEI10^oe^, notably in distal regions in females where CO frequency is low in wild-type and high in the mutants. In contrast, regions in the periphery of the centromeres where CO frequencies are relatively high in wild-type meiosis, correspond to recalcitrant zones with no or limited increase in the mutants. Zooming on closer centromere-proximal region, we examined the Non-Recombining Zones (NRZs) previously defined as the interval spanning the centromere with a complete absence of CO in the wild type, as scored in 3,613 gametes containing a total of 14,397 COs ^37^ (**Fig. 4d**). Strikingly, with such an unprecedented increase in CO numbers in the mutants described here, the NRZs persisted in resisting recombination and barely had any CO, despite our dataset containing a total of 70,022 COs. No COs were detected in NRZs in *zyp1 recq4* (10,825 COs), and only one CO was found in each population of *recq4* (3,675 COs), *recq4 figl1* (15,543 COs), and *zyp1 recq4* HEI10^oe^ (10,991 COs) (**Fig. 6d**). This strongly suggests that other mechanisms, that do not depend on HEI10, ZYP1, RECQ4 or FIGL1, prevent CO formation in the NRZs.

**Figure 4.**
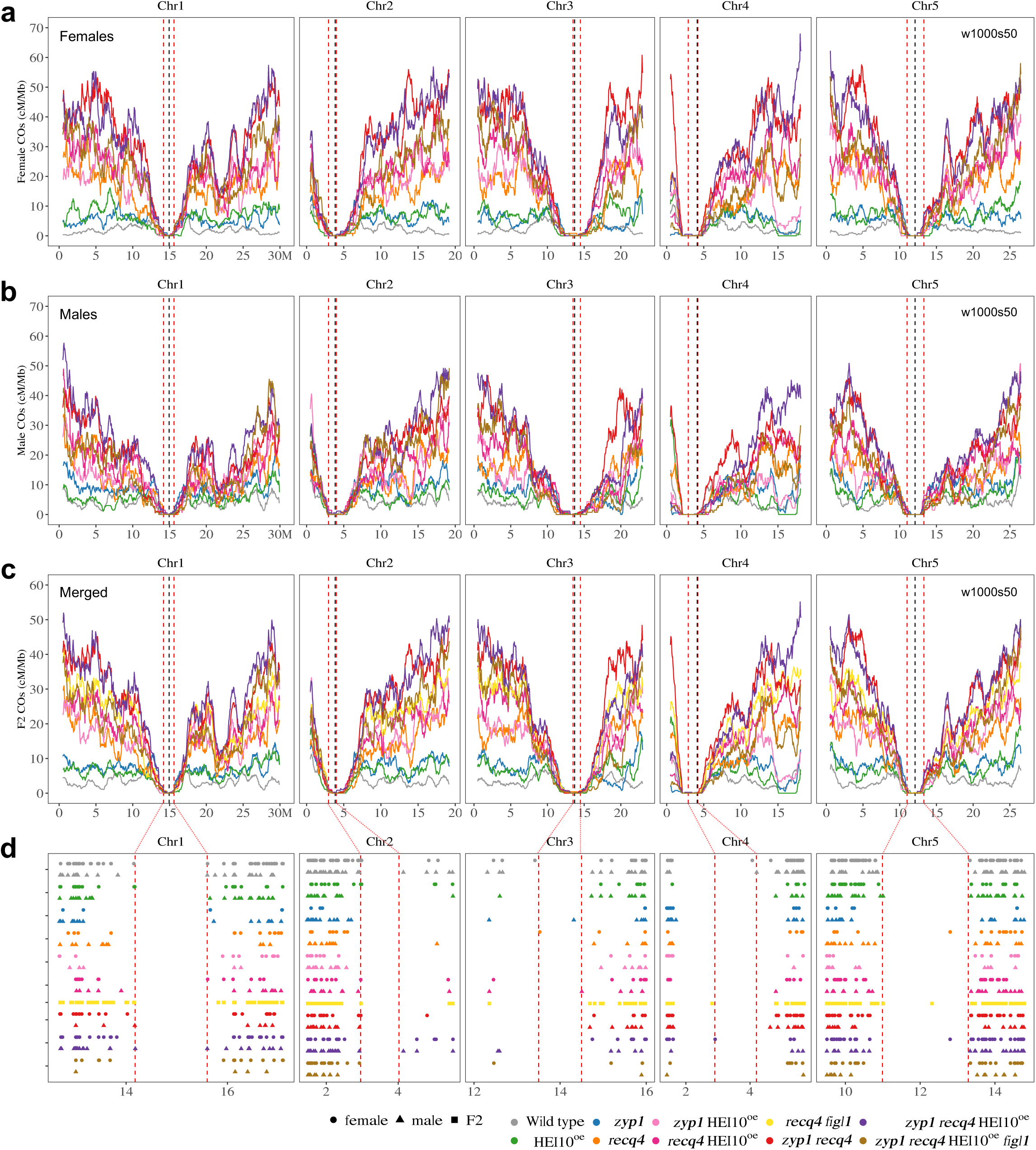
Chromosomal distribution of COs in female, male, and F2 contexts. a-b. The distribution (sliding window-based, window size 1 Mb, step size 50 kb) of COs along chromosomes in female (**a**) and male (**b**) of wild type, HEI10^oe^, *zyp1*, *recq4*, *zyp1* HEI10^oe^, *recq4* HEI10^oe^, *zyp1 recq4*, *zyp1 recq4* HEI10^oe^ and *zyp1 recq4* HEI10^oe^ *figl1*. **c.** The distribution (sliding window-based, window size 1 Mb, step size 50 kb) of COs along chromosomes in F2 or pseudo F2 of wild type, HEI10^oe^, *zyp1*, *recq4*, *zyp1* HEI10^oe^, *recq4* HEI10^oe^, *recq4 figl1*, *zyp1 recq4*, *zyp1 recq4* HEI10^oe^ and *zyp1 recq4* HEI10^oe^ *figl1*. **d.** The zoom of the CO position in the centromere proximal regions (Non-Recombining Zones, NRZs). Each point is a CO, circles, triangles and squares are females, males and F2s, respectively. The vertical dashed lines in red indicates the position of marker COs of NRZs, lines in black shows the middle position of centromeres.

**Figure 5.**
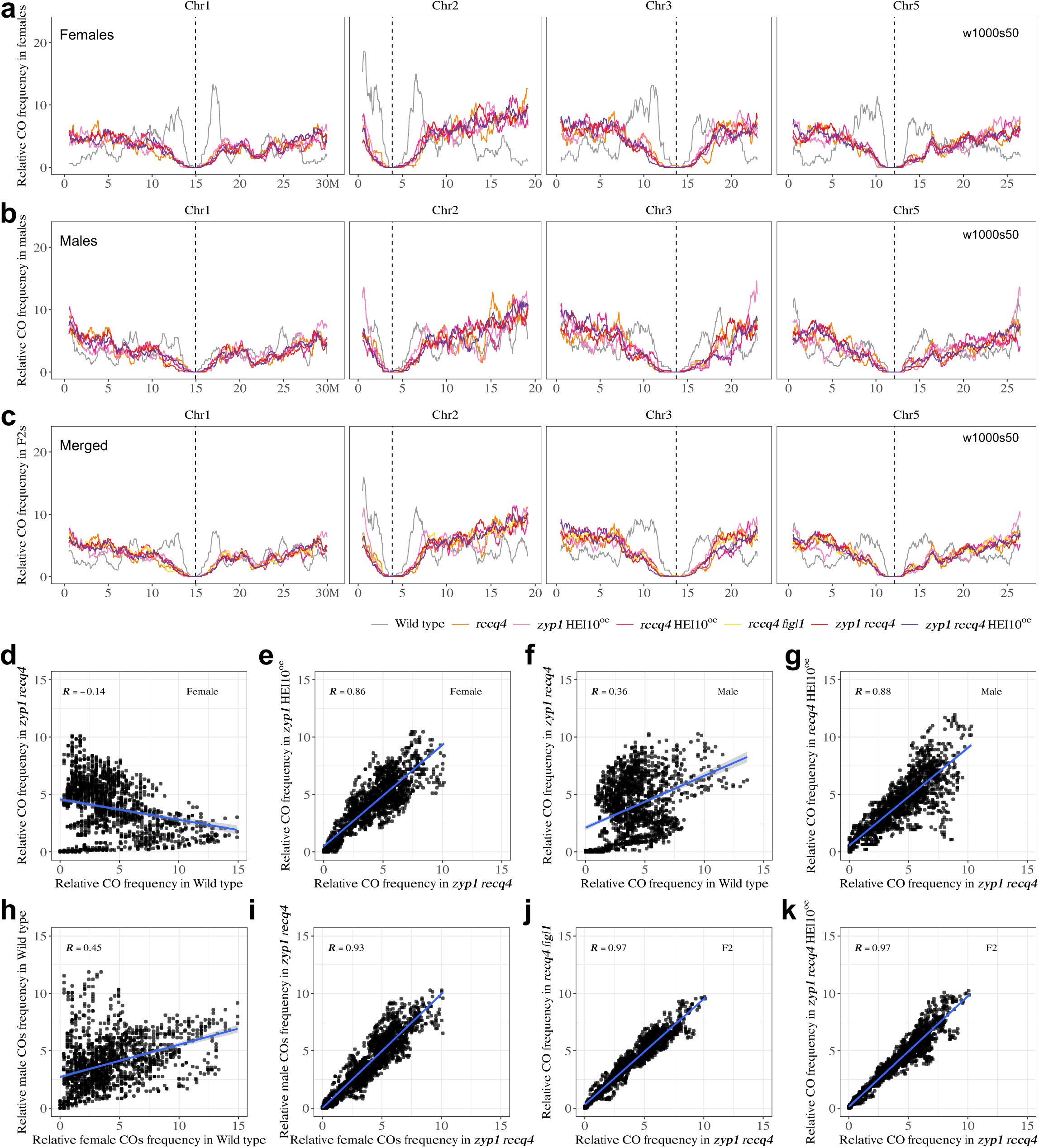
Comparison of the genome-wide CO landscape in female, male, and F2 contexts. a-b. The distribution (scaled per chromosome, sliding window-based, window size 1 Mb, step size 50 kb) of relative CO frequency along chromosomes in females (**a**) and males (**b**) of wild type, *recq4*, *zyp1* HEI10^oe^, *recq4* HEI10^oe^, *zyp1 recq4* and *zyp1 recq4* HEI10^oe^. **c.** The distribution (scaled per chromosome, sliding window-based, window size 1 Mb, step size 50 kb) of COs along chromosomes in F2 or pseudo F2 of wild type, *recq4*, *zyp1* HEI10^oe^, *recq4* HEI10^oe^, *recq4 figl1*, *zyp1 recq4* and *zyp1 recq4* HEI10^oe^. **d-k.** Spearman’s correlation tests of relative CO frequencies comparing between genotypes and sexes.

**Figure 6.**
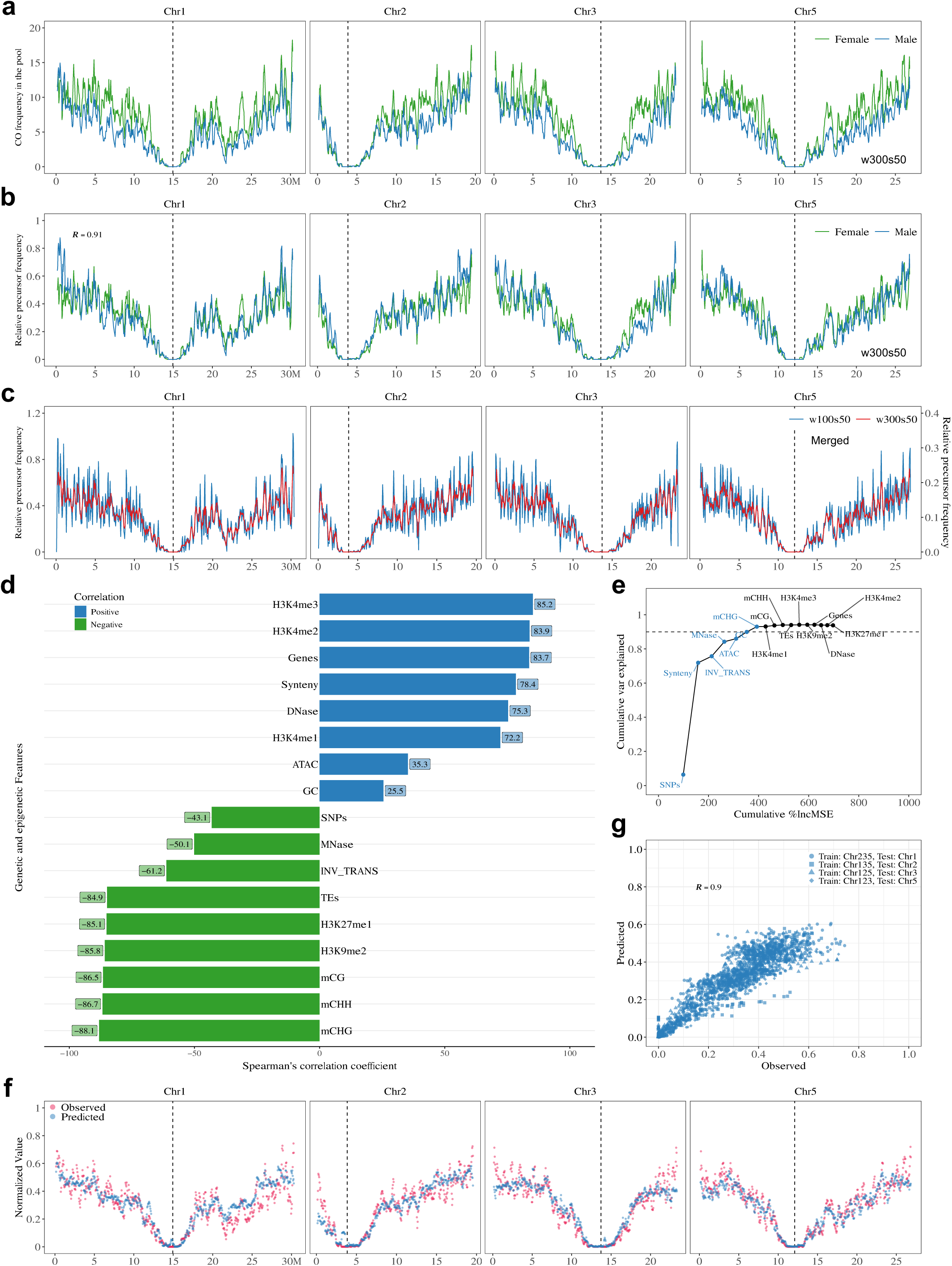
Association and prediction of precursor distribution with genetic and epigenetic features. **a.** The distribution (sliding window-based, window size 300 kb, step size 50 kb) of CO frequency along chromosomes in females and males, respectively, by merging *recq4*, *zyp1* HEI10^oe^, *recq4* HEI10^oe^, *zyp1 recq4*, *zyp1 recq4* HEI10^oe^ and *zyp1 recq4* HEI10^oe^ *figl1*. **b.** The distribution (scaled per genome, sliding window-based, window size 300 kb, step size 50 kb) of relative precursor frequency along chromosomes in females and males, respectively, by merging *recq4*, *zyp1* HEI10^oe^, *recq4* HEI10^oe^, *zyp1 recq4*, *zyp1 recq4* HEI10^oe^ and *zyp1 recq4* HEI10^oe^ *figl1*. **c.** The distribution (scaled per genome, sliding window-based, window size 100 kb and 300 kb, step size 50 kb) of relative precursor frequency along chromosomes in F2s, by merging females and males of *recq4*, *zyp1* HEI10^oe^, *recq4* HEI10^oe^, *zyp1 recq4*, *zyp1 recq4* HEI10^oe^ and *zyp1 recq4* HEI10^oe^ *figl1*, and F2s of *recq4 figl1*. The vertical dashed lines in black shows the middle position of centromeres. **d.** Spearman’s correlation test shows the comparison with features along chromosomes, with differences in colour and length according to the correlation scale. SNPs (SNPs density between Col and L*er*), INV_TRANS (inversions and translocations between Col and L*er*), Synteny (collinearity between Col and L*er*), Genes, TEs and GC (expressed gene in meiocytes, TE and GC density), ATAC and DNase (chromatin accessibility, ATAC-seq and DNase-seq, log2(Tn5/gDNA) and log2(DNase/gDNA), in floral tissues), H3K4me1/2/3, H3K9me2, H3K27me1 (euchromatin, heterochromatin, and Polycomb histone marks, ChIP-seq, log2(ChIP/input), in flower buds), mCG, mCHG and mCHH (DNA methylation in CG, CHG, and CHH contexts, proportion methylated cytosine, in male meiocytes), MNase (nucleosome occupancy, MNase-seq, log2(MNase/gDNA), in buds). **e.** The cumulated proportion of variation that can be explained with the features at the genome scale. The top seven most important features are coloured, for which the cumulative proportion of variation that can be explained reaches the plateau. **f.** The chromosomal distribution of observed and predicted precursor maps. The precursor profiles of individual chromosomes were predicted using profiles of the top seven most important features from the other three chromosomes. **g.** The Spearman’s correlation test between the predicted and observed precursor distributions. The training-testing dataset is differentiated in shapes.

Crossover landscapes in various mutants appear to have similar landscapes (**Fig. 4, 5**). To explore this formally, and compare the shape of the distributions independently of the absolute frequencies, we normalized the data by calculating the distribution of relative frequency per chromosome (i.e. what proportion of COs that occurred in this chromosome occurred in a given interval) (**Fig. 5 and Fig. S8-10**). In wild-type females, the density of COs was the highest in the periphery of centromeres, and low at chromosome ends (**Fig. 5a**). In the wild-type males, COs are also frequent in the periphery of centromeres, but chromosome ends are also CO-rich (**Fig. 5b**). Consequently, female and male distributions are poorly correlated in the wild type (Spearman’s correlation r =0.45, **Fig. 5h**). Both female and male *zyp1 recq4* distributions differ markedly from the wild-type distributions (**Fig. 5a, b**, R=-0.14 and R=0.36, **Fig. 5d, f**), but are strikingly similar to each other (r=0.93, **Fig. 5a, b, i**). Further, the CO distributions in various mutants were similar to each other, but distinct from the wild-type, in both females (e.g. r = 0.86, *zyp1* HEI10^oe^ vs *zyp1 recq4*; **Fig. 5a, d-e, Fig. S8**) and males (e.g. r = 0.88, *recq4* HEI10^oe^ vs *zyp1 recq4*; **Fig. 5b, f-g, Fig. S9**). The CO landscape in the F2 populations of *recq4 figl1* and *zyp1 recq4* are also strikingly consistent (r = 0.97, *recq4 figl1* vs *zyp1 recq4*; r = 0.97, *zyp1 recq4* HEI10^oe^ vs *zyp1 recq4*; **Fig. 5c, j-k, Fig. S10**) ^25^. CO landscapes in *recq4, zyp1* HEI10^oe^*, recq4* HE10^oe^*, recq4 figl1, zyp1 recq4, zyp1 recq4* HEI10^oe^ very all similar, with correlations in the range 0.75-0.92 (**Fig. S10**).

The convergence towards a similar pattern in females and males in different mutants suggests the existence of a common underlying feature. We propose that this distribution of Crossover Potential (CO_P_) represents the distribution of eligible recombination intermediates (i.e. DSBs that are being repaired with the homologous chromosomes, potentially leading to a CO). We then combined the 21,104 female and 15,188 male COs from the various mutant combinations (*recq4*, *zyp1* HEI10^oe^, *recq4* HEI10^oe^, *zyp1 recq4*, *zyp1 recq4* HEI10^oe^, and *zyp1 recq4* HEI10^oe^ *figl1*) to create high-resolution maps of CO_P_ in females and males, separately (**Fig. 6a**). The female and male CO_P_ maps are strikingly similar, the female curve being slightly above the male one. When normalized, the two CO_P_ curves almost perfectly overlap (**Fig. 6b**, Spearman’s correlation r = 0.91), showing that the distribution of CO_P_ is shared between the two sexes. These features of CO_P_ strikingly deviate from those of wild-type COs, which are more frequent in males than females and very differently distributed. We then combined the female and male COs of the mutants listed above, together with the *recq4 figl1* F2 population, totaling 49,482 COs, and generated a high-resolution universal CO_P_ map (**Fig. 6c**). The CO_P_ is not homogenous along chromosomes with null values in the centromeric regions and globally higher frequencies toward chromosome ends, and sharp peaks and valleys.

### Genomic features can predict the CO potential

We next wondered what could shape the CO_P_ landscape. We performed correlation analysis with 17 different genomic features (**Fig. 6d**), including GC content, meiotically expressed genes ^38^, transposable elements, sequence divergence between the two parental lines, chromatin accessibility, euchromatic and heterochromatic histone modification marks, DNA methylation, and nucleosome occupancy ^32^. The distribution of CO_P_ is strongly positively correlated with euchromatic histone modification markers (H3K4me3, r = 0.85), density of expressed genes in meiocytes (r = 0.84), open chromatin (DNase, r = 0.75), and synteny (r = 0.78). Oppositely, significant negative correlations were found with DNA methylation (mCHG, r = −0.88), heterochromatic histone modification marks (H3K9me2, r = −0.86), density of TEs (r = −0.85) and inversions and translocations (r = −0.61). This result suggests that CO_P_ is favored by certain open chromatin states and disfavored by sequence divergence.

We then employed a machine-learning algorithm (random forest) to measure the capacity of the 17 features to predict the CO_P_ landscape. Overall, 94% of the genome-wide variation can be explained by a model developed with all 17 features (**Fig. 6e**). Among the 17 features, SNP density was assessed of the utmost importance, which alone explained 6.4% of the genome-wide variation (**Fig. 6e**) and 28.4% of the variation along chromosome arms (**Fig. S11b**). By adding features step by step in the order of importance, we found that the top seven and eight features can explain 93% of the genome-wide variation and 90.6% of the variation along chromosome arms, respectively (**Fig. 6d, Fig. S11b**). To explore the predictive performance in the CO_P_ landscape of the model, we next performed 4 times cross-validation by using three chromosomes as the training dataset and one chromosome as the testing dataset (**Fig. 6f-g, Fig. S11c-d**). Considering the top seven and eight features for genome-wide and chromosome arms, the model trained from the training dataset worked well with the testing dataset, resulting in a strong correlation (r = 0.9, **Fig. 6f-g**, and r = 0.86, **Fig. S11c-d**) between the prediction and observation. The capacity of these features to accurately predict the landscape suggests that the CO_P_ might be entirely determined by genomic characteristics including chromatin state and sequence divergence.

## Discussion

Meiotic crossover frequencies are naturally limited to a few per chromosome. Here, we have shown an unprecedented elevation of meiotic crossovers in Arabidopsis by simultaneously mutating the synaptonemal complex ZYP1 and the anti-recombination helicase RECQ4. We observed an average of 34.7 and 23.5 COs per female and male *zyp1 recq4* gametes, respectively, corresponding to 58 COs per generation, to be compared with 8.1 in wild-type and 50 in the previous champion *recq4 figl1* ^3,25^. The fertility is only moderately affected in *zyp1 recq4* and no genomic instability was observed, opening a novel possibility to manipulate recombination for the benefit of plant breeding. Higher CO levels could be leveraged to enhance genetic mixing genetic information in the early steps of breeding, reduce the size of introgression to elite lines, and facilitate the identification of genetic determinants of traits ^25^. *RECQ4* mutation increases CO numbers in rice, tomato, and pea ^39,40^, suggesting that manipulation of this factor can be useful in a wide variety of crops. Mutation of *zyp1* in rice also increased class I CO number, but a *ZYP1* RNAi reduced them in Barley ^41,42^, calling for testing the effects of *zyp1*, and *zyp1 recq4* in more species.

Mutating both ZYP1 and RECQ4 provoked a remarkable elevation in CO numbers, beyond what was expected under the hypothesis of simple additive effects of both mutations on class I and class II COs, respectively. These massive increases correspond to class II COs, as indicated by MLH1 not marking extra COs. This points to a function of ZYP1 in preventing class II COs, which is revealed only when RECQ4 is absent, in addition to its documented function in regulating class I COs ^17,18^. It suggests that recombination precursors (DNA double-strand breaks) could be increased in the absence of ZYP1, as shown when the homologous protein *Zip1* is defective in yeast ^43,44^. If the same scenario happens in *zyp1 recq4*, the additional DSBs would produce additional recombination intermediates, that would be repaired as non-crossover in the presence of RECQ4, but converted into class II COs in its absence. Another non-exclusive possibility is a role of the synaptonemal complex in preventing the formation of aberrant recombination intermediates (e.g. Multiple invasions from a single DSB). This is supported by the slight chromosome fragmentation observed in *zyp1 mus81*, pointing to the role of the nuclease MUS81 in repairing recombination intermediated in the absence of ZYP1. The more extensive fragmentation observed in rice *zyp1 mus81* ^45^ suggests that this function could be more prominent in this species. Finally, one may speculate a role of ZYP1 in channeling the repair of a proportion of DSBs to the sister chromatid, leading to an increase of inter-homolog intermediate in *zyp1* that would synergize with the anti-CO effect of RECQ4.

Combining HEI10^oe^*, zyp1* or *recq4* two by two systematically increased CO beyond the level of the single mutants. This predicted that combining the three should increase even further CO frequencies. However, adding HEI10^oe^ to the double with the strongest effect *zyp1 recq4* did not further increase COs. Interestingly, HEI10^oe^ did increase the number of MLH1 foci in both sexes in *zyp1 recq4*, without changing the total number of COs. This suggests that overexpressing HEI10 does, as expected, increase the number of class I COs, but that this is at the expense of class II COs. This suggests that a maximum number of COs has been reached and that at these high levels, the class I and class II pathways compete for a limited number of precursors. Intriguingly, while *figl1* can enhance CO frequencies in *recq4* ^3,25^, adding the *figl1* mutation to *zyp1 recq4* HEI10^oe^ does not increase COs, but decreases the CO frequency. We suspect that in this context, aberrant recombination molecules are formed, limiting their possibility to be matured into COs. In any case, it appears that the CO frequency observed in *zyp1 recq4* corresponds to some kind of maximum. This maximum corresponds to ∼70 chiasmata/COs per female meiocyte (average of 35 COs per gamete) and ∼48 chiasmata/COs per male meiocyte (average of 24 COs per gamete). Based on DMC1/RAD51 foci, it is estimated that ∼200 DSBs are produced per male meiocyte ^46^. If we consider that 1/3 are repaired on the sister (absence of bias, one sister chromatid and two homologous chromatids) ^47^, and that the class II pathway is not biased in the repair of CO precursors (1:1 CO/NCO), we would expect ∼66 COs, to compare with the estimated ∼48 chiasmata in males. If all the considerations are correct, this suggests that additional mechanisms prevent DSB maturation into COs. Intriguingly, we observed an inversion of heterochiasmy in all mutants in which class II COs is deregulated, with more COs in females than in males. One parsimonious hypothesis is that in all contexts, including wild-type, more DSBs are formed in females than males, but that the CO designation process in wild-type which is linked to the length of the SC and impassive to DSB counts would lead to fewer COs in females than males.

Strikingly, in female and male meiosis of various mutants, the CO distribution converged to a common profile leading us to propose the concept of Crossover Potential (CO_P_) whose density map is revealed when the CO designation process is deregulated. In this view, the CO_P_ determines where COs might occur, combining the local capacity to experience DNA double-strand breaks and the availability of a viable repair template on the homologous chromosome. Accordingly, open chromatin markers (positively correlated) and sequence divergences (negatively correlated) can together efficiently predict CO_P_. CO_P_ would be responsible for the local placement of COs in genes observed in many species ^48–52^. Note that PRDM9 which drives DSB positions in many mammals dramatically affects CO_P_ distribution ^53^. Importantly, CO_P_ very partially dictates the final CO distribution because of the CO designation process ^54,55^ that determines the fate of eligible recombination intermediates. A striking example is the sexual dimorphism in the hermaphrodite Arabidopsis, where female and male CO landscapes markedly differ despite an identical genome and CO_P_ map. The limited influence of CO_P_ on the global CO distribution explains why the megabase-scale CO landscape is largely independent of sequence divergence ^32^. While CO_P_ appears to be determined at the genomic level (chromatin state and sequence divergence), the CO designation process is a chromosomal event influenced by the higher-order spatial organization in the synaptonemal complex ^20,56,57^. The CO distribution is thus determined by the combination of the CO_P,_ which is particularly important at the local scale and can dismiss some regions, and the designation process that shapes the global chromosomal landscape and dictates CO counts.

## Material and Methods

### Plant materials and growth conditions

*Arabidopsis thaliana* plants were cultivated in Polyklima growth chambers (16-h day, 21.5 °C, 280 µM; 8-h night, 18 °C: 60% humidity). The following Arabidopsis lines were used in this study: Wild type Col-0(186AV1B4) and L*er*−1 (213AV1B1) from the Versailles stock center (http://publiclines.versailles.inra.fr/). This study used *zyp1-1* in Col and *zyp1-6* in L*er* ^17^, *recq4a* in Col (*recq4a-4*, N419423) ^58^, *recq4a-W387X* in L*er* ^21^*, recq4b* in Col (*recq4b-2*, N511330) ^58^, *figl1-19* in Col (*SALK_089653*) and *figl1-12* in L*er* ^22^. The HEI10 over-expression line is Col HEI10 line C2 ^16^.

### Generation of multi-mutants

The combinations of different mutants (Col/L*er* hybrids) were obtained by a series of crossing schemes (Table S1 and Fig. S1-4). *zyp1* HEI10^oe^ was produced previously^20^. These eight different mutants and their sister wild-type control plants were backcrossed with Col male and female plants, respectively, to get the BC1 population for sequencing and subsequent CO analysis. BC1 populations were grown for three weeks (16-h day/ 8-h night) and four days in the dark. For DNA extraction and library preparation, 100–150 mg leaf samples were collected from the various BC1 populations ^59^.

### CO identification and analysis

In this study, the female and male population of wild type (95 and 89 plants), HEI10^oe^ (48 and 46 plants), *zyp1* (42 males), *recq4* HEI10^oe^ (142 and 139 plants), *zyp1 recq4* (186 and 186 plants), *zyp1 recq4* HEI10^oe^ (190 and 177 plants), *zyp1 recq4* HEI10^oe^ *figl1* (138 and 134 plants), were sequenced by Illumina HiSeq3000 (2×150bp) conducted by the Max Planck-Genome-center (https://mpgc.mpipz.mpg.de/home/). The raw sequencing data of the female and male population of wild type (428 and 294 plants, ArrayExpress number: E-MTAB-11254 ^32^), HEI10^oe^ (144 and 141 plants, ArrayExpress number: E-MTAB-11696 ^20^ and *zyp1* (272 and 225 plants, ArrayExpress number: E-MTAB-9593 ^17^, E-MTAB-11696 ^20^) from previous studies were also included in this study. In total, we analyzed female and male populations of 523 and 383 wild type, 192 and 187 HEI10^oe^, 272 and 267 *zyp1*, 142 and 138 *recq4*, 117 and 116 *zyp1* HEI10^oe^, 142 and 139 *recq4* HEI10^oe^, 186 and 186 *zyp1 recq4*, 190 and 177 *zyp1 recq4* HEI10^oe^, 138 and 134 *zyp1 recq4* HEI10^oe^ *figl1* plants.

The raw sequencing data were quality-controlled using FastQC v0.11.9 (http://www.bioinformatics.babraham.ac.uk/projects/fastqc/). The sequencing reads were aligned to the *Arabidopsis thaliana* Col-0 TAIR10 reference genome ^60,61^, using BWA v0.7.15-r1140 ^62^, with default parameters. A set of Sambamba v0.6.8 ^63^ commands was used for sorting and removing duplicated mapped reads. The creation of the high-confidence SNP marker list between Col and L*er*, meiotic CO detection (a sliding window-based method, with a window size of 30 kb and a step size of 15 kb), check and filtering of low covered and potential contaminated samples were performed according to previous protocols ^17,20,32,64,65^. Samples of each population were randomly selected to check predicted COs manually by inGAP-family ^65^.

The Coefficient of Coincidence (CoC) was calculated using MADpattern ^57,66^, with 10 intervals. The chromosome 4 was excluded from interference analyses because of a translocation associated with the HEI10 transgene ^15^ and potential inversion in our L*er* line ^17^.

For CO distribution analysis, we refined the position of the marker COs of each NRZ boundary against the Col-TAIR10 reference genome used in this study. In addition, it should be noted that a translocation associated with the HEI10 C2 transgene at the short arm and a megabase-scale inversion at the long arm of chromosome 4, which may introduce bias of the CO distribution and thus chromosome 4 was excluded ^15^.

To define hot and cold zones of CO burst from hyper recombination mutants, we examined COs in 1-Mb windows with 50-kb sliding and defined (i) the hot zones as common regions in *zyp1* HEI10^oe^ and at least 3 other mutants, with at least a two-fold increase than the median compared with wild-type, and (ii) the cold zones as common regions in *zyp1* HEI10^oe^ and at least 3 other mutants, with at most a half-fold increase than the median compared with wild-type.

### Aneuploidy screening by whole–genome sequencing

The genome was first cut into non-overlapping 100 kb windows, whose sequencing depth was estimated by Mosdepth v0.2.7^67^ with parameters of “-n–fast-mode–by 100000”. Then, pairwise testing of sequencing depths between chromosomes of individuals was performed using the Mann–Whitney test, and the p values were adjusted using the fdr method. A pair of tested chromosomes with fold change >1.2 and p value <1e−20 was considered as aneuploid.

### Meiocyte RNA-seq analysis

The RNA-seq dataset of Arabidopsis thaliana meiocytes from a previous study ^38^ was downloaded from the NCBI SRA database (SRR5209212 and SRR5209213). First, quality checking of the raw sequencing reads was performed by using FastQC. Then, HISAT2 v2.1.0^68^ was used to align the reads against the Col-0 TAIR10 reference genome. Gene expression was normalized as TPM, which was calculated by StringTie v2.0.6 69 with default parameters.

### Cytology and Image processing

Meiotic chromosome spreads were performed as previously described^70^. Chromosomes were stained with DAPI (2μg/ml). Slides were observed using a Zeiss Axio Imager Z2 microscope. Images were acquired under a 100 × oil immersion objective, and processed with ZEN software.

Immunolocalization performed on cells with preserved three-dimensional structures on male and female meiocytes was performed as previously described ^17,20^. Three primary antibodies were used for MLH1 foci number counting: anti-REC8 raised in rat ^71^ (laboratory code PAK036, dilution 1:250), anti-MLH1 raised in rabbit ^72^ (PAK017, 1:200), and anti-HEI10 raised in chicken ^73^ (PAK046, 1:5,000). Images were acquired under Laica THUNDER Imager system and Abberior instrument facility line (https://abberior-instruments.com/). For Leica THUNDER Imager system, 555-nm, 635-nm and 475-nm excitation lasers were used for STAR Orange, STAR Red and STAR GREEN, respectively. Images were deconvoluted using Large Volume Computational Clearing (LVCC) mode in a Laica LAS X 5.1.0 software. For Abberior instrument facility line, images were acquired with 561- and 640-nm excitation lasers (for STAR Orange and STAR Red, respectively) and a 775-nm STED depletion laser. Confocal images were taken with the same instrument with a 485-nm excitation laser (for STAR GREEN/ Alexa488). Images were deconvoluted by Huygens Essential (version 21.10, Scientifific Volume Imaging, https://svi.nl/). Deconvoluted pictures were imported to Imaris x64 9.6.0 (https://imaris.oxinst.com/, Oxford Instruments, UK) for foci counting. The Spots module was used for MLH1 foci counting in diplotene and diakinesis cells. The majority of MLH foci colocalize with HEI10 foci. Only double MLH1/HEI10 foci present on chromosomes were taken into account.

## Supporting information

Supplemental data Table CO list

## Acknowledgments

This work was supported by core funding from the Max Planck Society, and Alexander von Humboldt Fellowships to Q.L. and J.J. We thank Ian Henderson for kindly providing the HEI10 C2 line.

## Author contributions

R.M. lead the project. J.J. produced the genetic material, analyzed the cytology experiments and fertility (wild type, *zyp1*, *recq4*, *zyp1 recq4*, *zyp1* HEI10^oe^, *recq4* HEI10^oe^, *zyp1 recq4* HEI10^oe^ and *zyp1 mus81*). Q.L. analyzed the sequencing data and performed recombination, interference, and aneuploidy analyses. S.D. developed the protocol for female immunolocalization, produced the genetic material and analyzed the cytology experiments and fertility (*zyp1 recq4* HEI10^oe^ *figl1*). J.J. Q.L. and R.M. wrote the manuscript.

## Data availability

The list of identified COs in the female and male populations of wild type, HEI10^oe^, *zyp1*, *recq4*, *zyp1* HEI10^oe^, *recq4* HEI10^oe^, *zyp1 recq4*, *zyp1 recq4* HEI10^oe^ and *zyp1 recq4* HEI10^oe^ *figl1* can be accessed in Supplementary Data 1. The raw sequencing data generated in this study have been deposited in the ArrayExpress EMBL-EBI database under accession codes E-MTAB-14424, E-MTAB-14425, E-MTAB-14426, E-MTAB-14427, E-MTAB-14428 and E-MTAB-14430. Source data are provided with this paper.

