## Supplemental figures for "Maximizing meiotic crossover rate reveals the map of Crossover Potential"

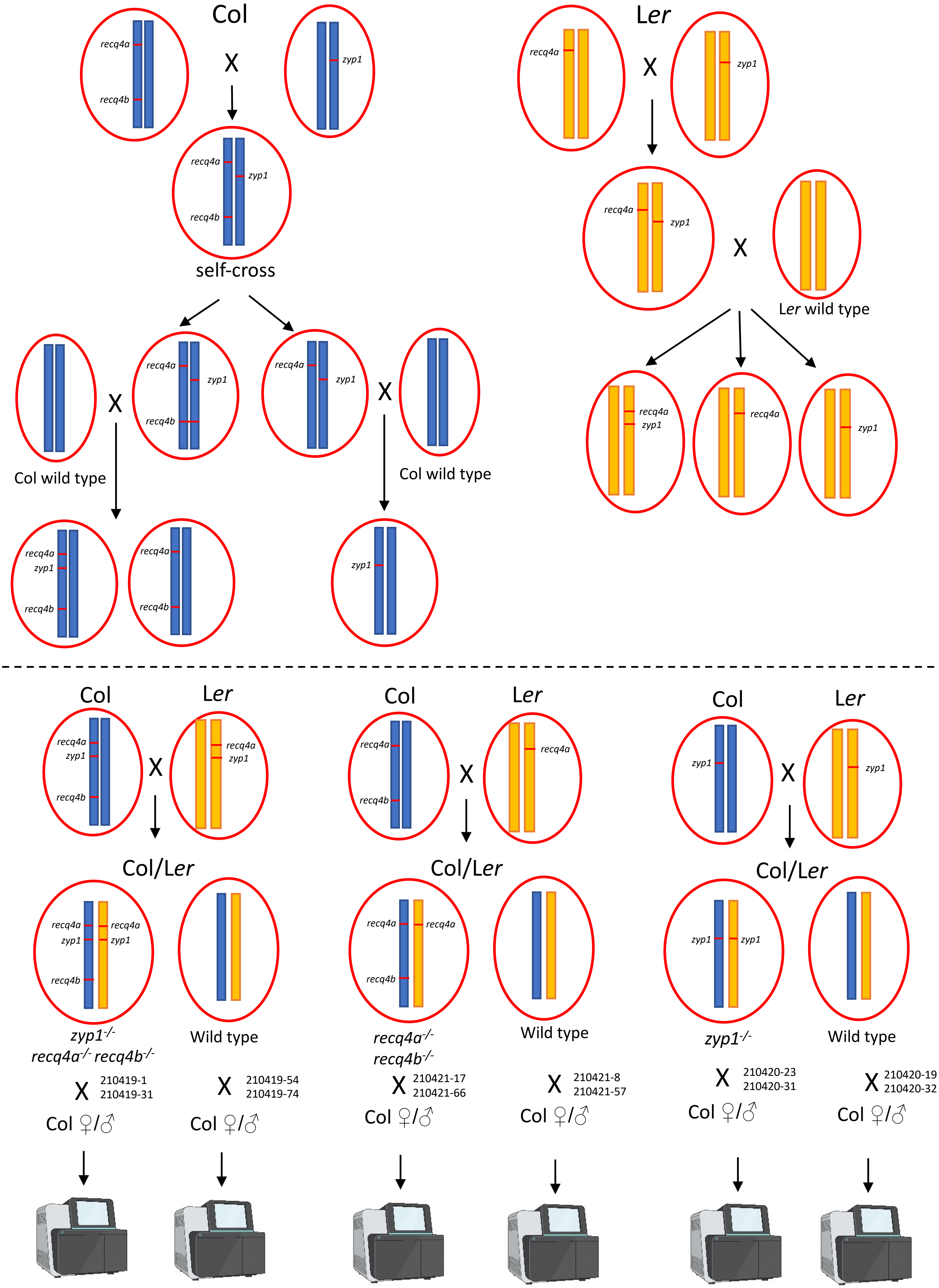

Col
X
recq4a
recq4b
zyp1
self-cross
Col wild type
Ler
Ler wild type
Col/Ler
zyp1-/- recq4a-/- recq4b-/-
recq4a-/- recq4b-/-
zyp1-/-
Wild type
210419-1
210419-31
210419-54
210419-74
210421-17
210421-66
210421-8
210421-57
210420-23
210420-31
210420-19
210420-32
Col ♀/♂

**Fig S1. Generation of *zyp1*$^{-/-}$ *recq4a*$^{-/-}$ *recq4b*$^{-/-}$(Col/L*er*), *recq4a*$^{-/-}$ *recq4b*$^{-/-}$ (Col/L*er*), *zyp1*$^{-/-}$ (Col/L*er*) and their respective wild type (Col/L*er*) controls.**

*zyp1*$^{+/-}$(Col) plants were crossed with *recq4a*$^{+/-}$ *recq4b*$^{+/-}$*(*Col) plants to get *zyp1*$^{+/-}$ *recq4a*$^{+/-}$ *recq4b*$^{+/-}$ (Col), and then self-crossed to get *zyp1*$^{+/-}$ *recq4a*$^{+/-}$ *recq4b*$^{-/-}$*(*Col) and *recq4a*$^{+/-}$ *zyp1*$^{+/-}$(Col) plants. Note that ZYP1, RECQ4A and RECQ4B are all located on chromosome 1, to make sure the mutation of the three genes are all on the same chromosome , *zyp1*$^{+/-}$ *recq4a*$^{+/-}$ *recq4b*$^{-/-}$(Col. *zyp1*, *recq4a*, *recq4b* mutation in trans) were back crossed with Col wild type to get *zyp1*$^{+/-}$ *recq4a*$^{+/-}$ *recq4b*$^{+/-}$*(*Col. zyp1, recq4a and recq4b in cis) and *recq4a*$^{+/-}$ *recq4b*$^{+/-}$(Col. *recq4a*, *recq4b* in cis); *recq4a*$^{+/-}$ *zyp1*$^{+/-}$ (Col, *recq4a* and *zyp1* in trans) were crossed with Col wild type to get zyp1$^{+/-}$(Col). *zyp1*$^{+/-}$(L*er*) were crossed with *recq4a*$^{+/-}$(L*er*) to get *zyp1*$^{+/-}$ *recq4a*$^{+/-}$(L*er, zyp1* and recq4 in trans), then *zyp1*$^{+/-}$ *recq4a*$^{+/-}$ (L*er*, *zyp1* and *recq4* in trans) were backcrossed with wild type (L*er*) to get *zyp1*$^{+/-}$ *recq4a*$^{+/-}$(L*er*, *zyp1* and *recq4* in cis), *recq4a*$^{+/-}$(L*er*) and *zyp1*$^{+/-}$(L*er*). Then *zyp1*$^{+/-}$ *recq4a*$^{+/-}$ *recq4b*$^{+/-}$ (Col, *zyp1*, *recq4a* and *recq4b* in cis) were crossed with *zyp1*$^{+/-}$ *recq4a*$^{+/-}$ (L*er*, *zyp1* and *recq4a* are in cis) to get zyp1$^{-/-}$ recq4a$^{-/-}$ recq4b$^{-/-}$(Col/L*er*) and sister wild type(Col/L*er*) control; *recq4a*$^{+/-}$ *recq4b*$^{+/-}$ (Col) were crossed with *recq4a*$^{+/-}$ (L*er*) to get *recq4a*$^{-/-}$ *recq4b*$^{-/-}$ (Col/L*er*) and sister wild type(Col/L*er*) control; *zyp1*$^{+/-}$(Col) were crossed with *zyp1*$^{+/-}$(L*er*) plants to get *zyp1*$^{-/-}$(Col/L*er*) and sister wild type(Col/L*er*) control.

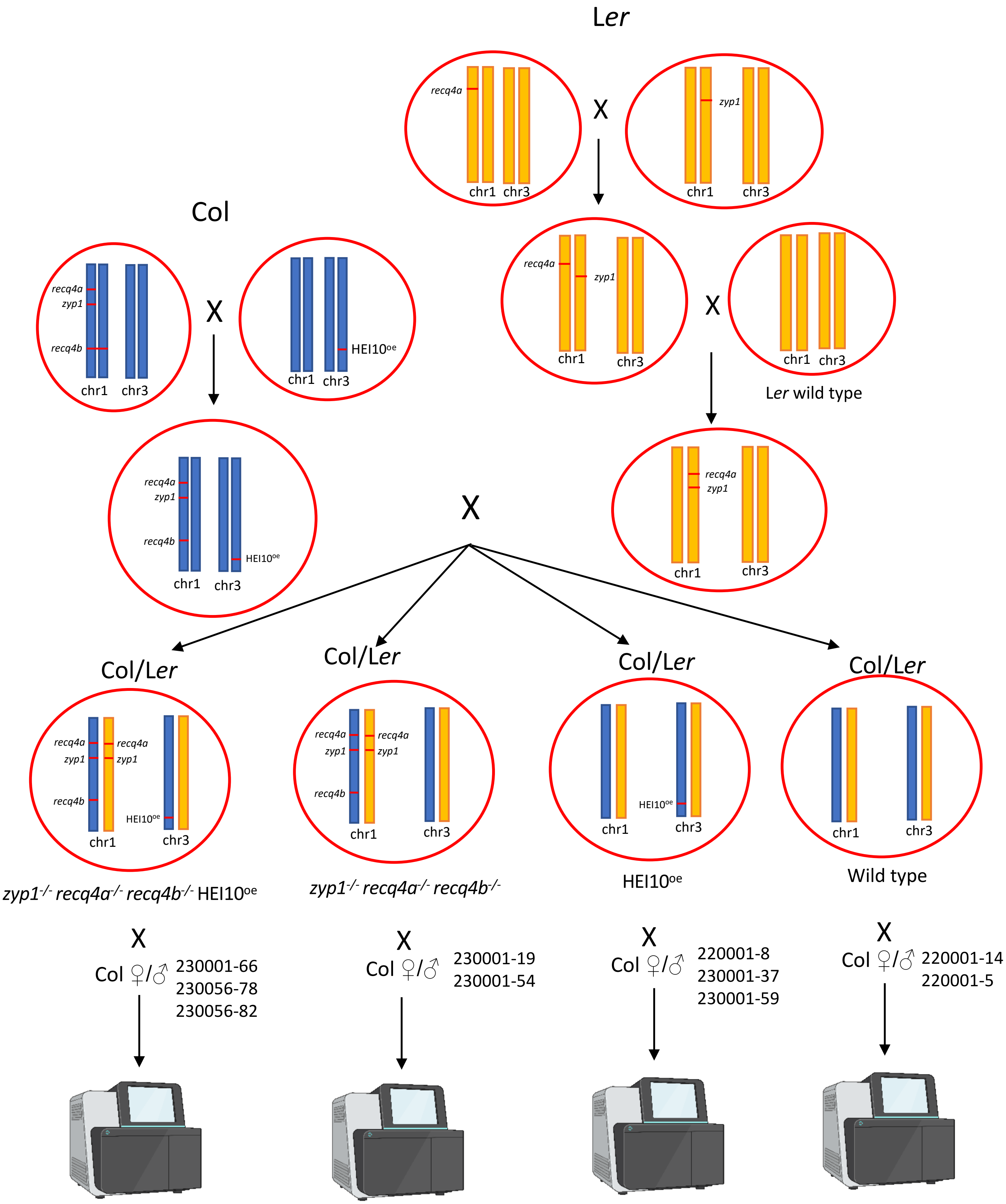


**Fig S2. Generation of *zyp1⁻/⁻ recq4a⁻/⁻ recq4b⁻/⁻* HEI10ᵒᵉ (Col/*Ler*), *zyp1⁻/⁻ recq4a⁻/⁻ recq4b⁻/⁻* (Col/*Ler*), HEI10ᵒᵉ (Col/*Ler*) and wild type (Col/*Ler*)**

*zyp1*$^{+/-}$ *recq4a*$^{+/-}$ *recq4b*$^{-/-}$ (Col) were crossed with HEI10oehz (Col) to get *zyp1*$^{+/-}$ *recq4a*$^{+/-}$ *recq4b*$^{+/-}$ HEI10oehz (Col. *zyp1*, *recq4a* and *recq4b* in cis), *then zyp1*$^{+/-}$ *recq4a*$^{+/-}$ *recq4b*$^{+/-}$ HEI10oehz (Col. *zyp1*, *recq4a* and *recq4b* in cis) were crossed with *zyp1*$^{+/-}$ *recq4a*$^{+/-}$(*Ler*, zyp1 and *recq4a* in cis) to get *zyp1*$^{-/-}$ *recq4a*$^{-/-}$ *recq4b*$^{-/-}$ HEI10oehz (Col/*Ler*), *zyp1*$^{-/-}$ *recq4a-/- recq4b*$^{-/-}$ (Col/*Ler*), HEI10oehz (Col/*Ler*) and wild type.

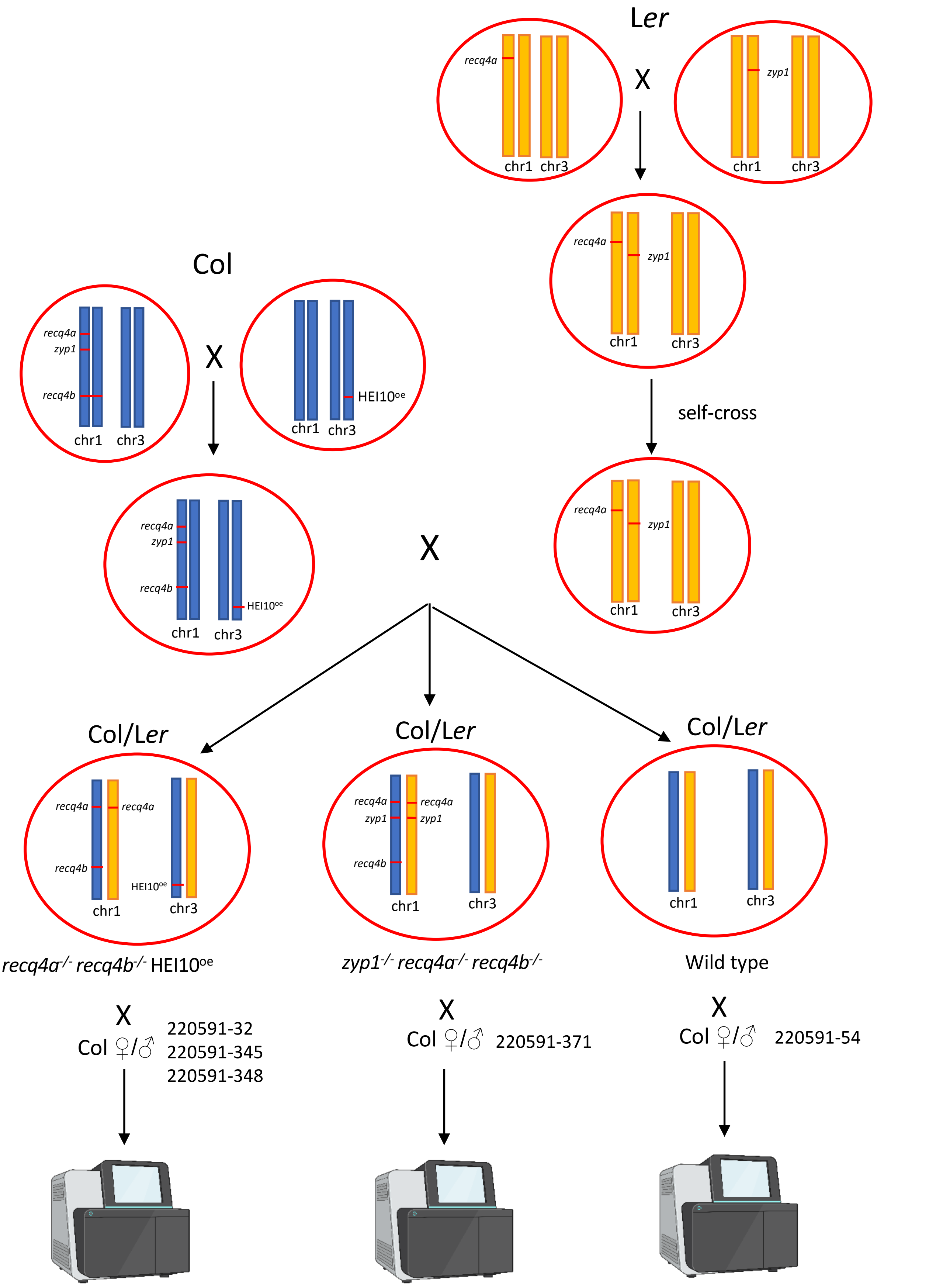


**Fig S3. Generation of *recq4a*$^{-/-}$ *recq4b*$^{-/-}$ HEI10$^{oe}$ (Col/*Ler*), *zyp1*$^{-/-}$ *recq4a*$^{-/}$ *recq4b*$^{-/-}$ (Col/*Ler*), wild type (Col/*Ler*).**

*zyp1*$^{+/-}$ *recq4a*$^{+/-}$ *recq4b*$^{-/-}$ (Col) were crossed with HEI10oehz (Col) to get *zyp1*$^{+/-}$ *recq4a*$^{+/-}$ *recq4b*$^{+/-}$ HEI10oehz (Col. *zyp1*, *recq4a* and *recq4* in cis), *zyp1*$^{+/-}$ *recq4a*$^{+/-}$ *recq4b*$^{+/-}$ HEI10oehz (Col, *zyp1*, *recq4a* and *recq4b* in cis) were crossed with *zyp1*$^{+/-}$ *recq4a*$^{+/-}$ (L*er*, *zyp1* and *recq4a* in trans) to get *recq4a*$^{-/-}$ *recq4b*$^{-/-}$ HEI10oehz (Col/L*er*), *zyp1*$^{-/-}$ *recq4a*$^{-/-}$ *recq4b*$^{-/-}$ (Col/L*er*) and wild type (Col/L*er*).

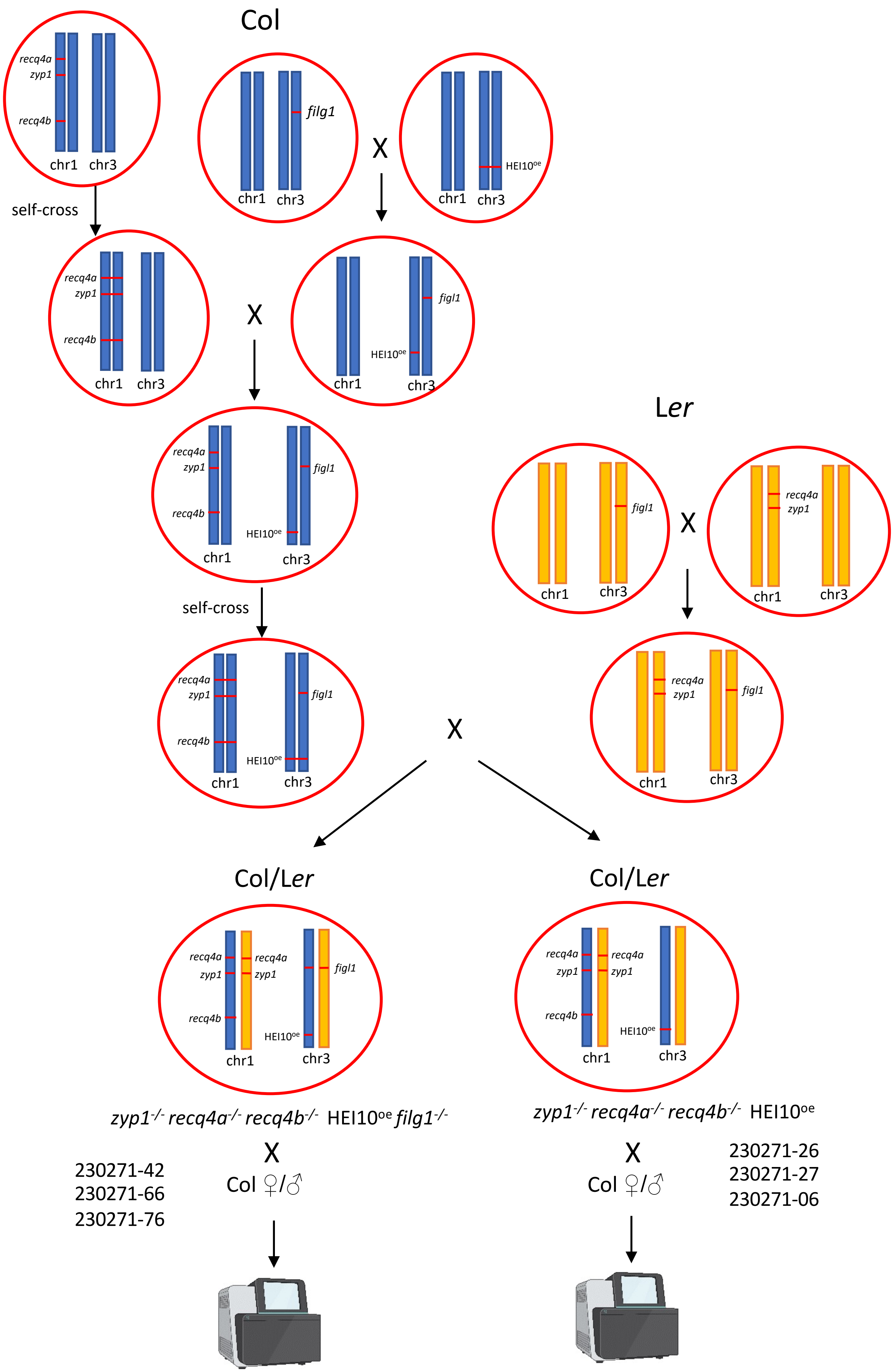

Col
recq4a
zyp1
recq4b
chr1
chr3
filg1
X
HEI10oe
self-cross
figl1
Ler
Col/Ler
zyp1-/- recq4a-/- recq4b-/- HEI10oe filg1-/-
zyp1-/- recq4a-/- recq4b-/- HEI10oe
Col ♀/♂
230271-42
230271-66
230271-76
230271-26
230271-27
230271-06

**Fig S4. Generation of *zyp1$^{-/-}$ recq4a$^{-/-}$ recq4b$^{-/-}$* HEI10$^{oe}$ *figl1$^{-/-}$* (Col/L*er*), and *zyp1$^{-/-}$ recq4a$^{-/-}$ recq4b$^{-/-}$* HEI10$^{oe}$ (Col/L*er*).**

*zyp1$^{+/-}$ recq4a$^{+/-}$ recq4b$^{+/-}$*(Col. *zyp1, recq4a* and *recq4b* in cis) were self-crossed to get *zyp1$^{-/-}$ recq4a$^{-/-}$ recq4b$^{-/-}$* (Col), *figl1$^{+/-}$*(Col) were crossed with HEI10oe homo (Col) to get HEI10oehz *figl1$^{+/-}$* (Col). Then *zyp1-/- recq4a-/- recq4b$^{-/-}$* (Col) were crossed with HEI10oehz *figl1$^{+/-}$*(Col, HEI10oehz and *figl1* in trans) to get *zyp1$^{+/-}$ recq4a$^{+/-}$ recq4b$^{+/-}$* HEI10oehz *figl1$^{+/-}$*(Col. *zyp1, recq4a* and *recq4b* in cis, HEI10oehz and *figl1* in trans we don't know), then this plant was self-crossed to get *zyp1-/- recq4a-/- recq4b-/-* HEI10oehomo *figl1$^{+/-}$* (Col). *zyp1$^{+/-}$ recq4a$^{+/-}$* (L*er*, *zyp1* and *recq4a* in cis) were crossed with *figl1$^{+/-}$*(L*er*) to get *zyp1$^{+/-}$ recq4a$^{+/-}$ figl1$^{+/-}$* (L*er*, *zyp1* and *recq4a* in cis). Then *zyp1$^{-/-}$ recq4a$^{-/-}$ recq4b$^{-/-}$* HEI10oehomo *figl1$^{+/-}$* (Col) were crossed with *zyp1$^{+/-}$ recq4a$^{+/-}$ figl1$^{+/-}$* (L*er*, zyp1 recq4a in cis) to get *zyp1$^{-/-}$ recq4a$^{-/-}$ recq4b$^{-/-}$* HEI10oehz *figl1$^{-/-}$* (Col/L*er*) and *zyp1$^{-/-}$ recq4a$^{-/-}$ recq4b$^{-/-}$* HEI10oe hz (Col/L*er*) (Fig. S4).

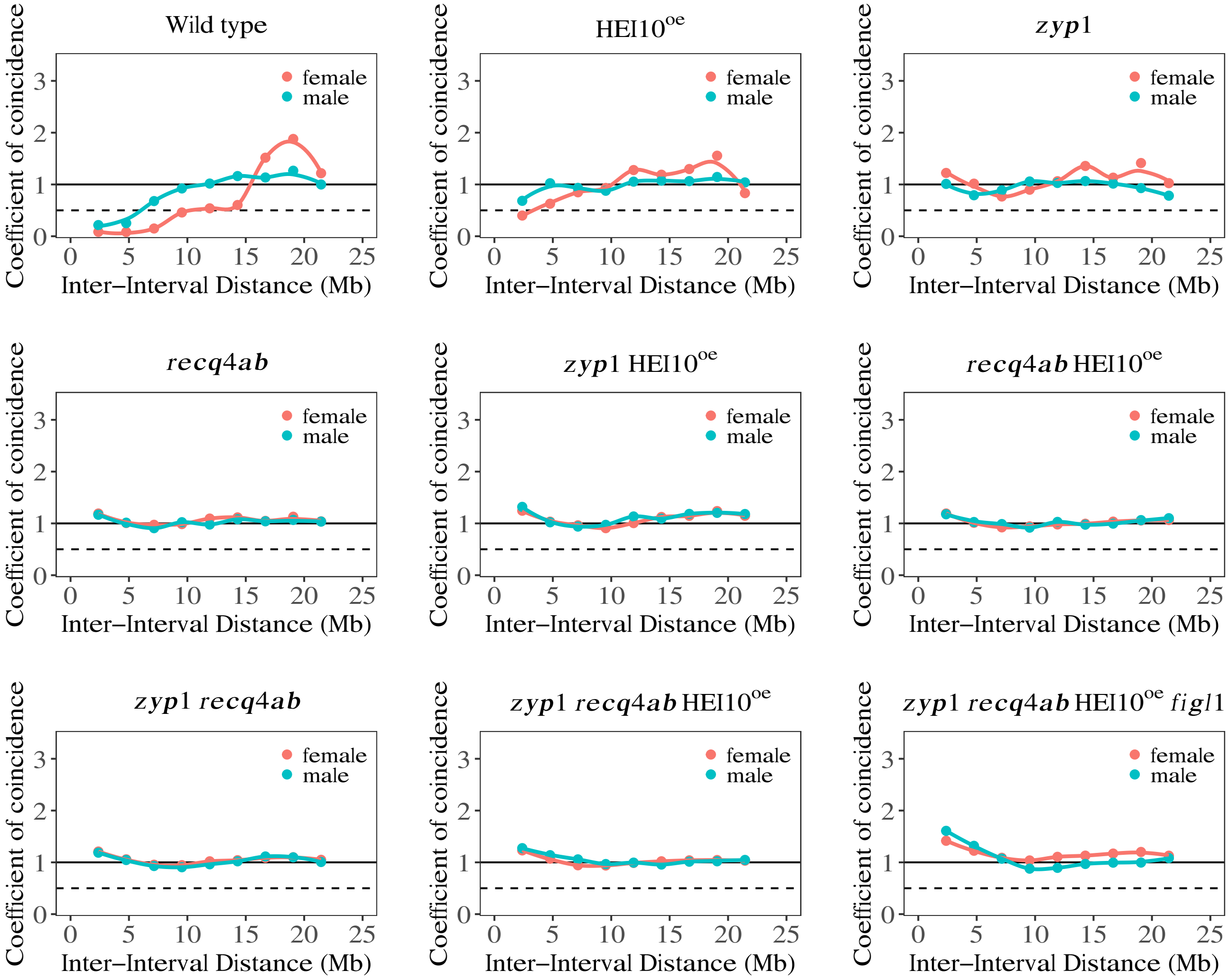


**Fig S5. Analysis of CO interference.**

CoC curves in the female and male meiosis of wild type, HEI10$^{oe}$, *zyp1*, *recq4*, *zyp1* HEI10$^{oe}$, *recq4* HEI10$^{oe}$, *zyp1 recq4*, *zyp1 recq4* HEI10$^{oe}$ and *zyp1 recq4* HEI10$^{oe}$ *figl1*, respectively. Chromosomes were divided into 10 intervals, for calculating the mean coefficient of coincidence of each pair of intervals. A CoC close to 1 means the absence of CO interference. A CoC close to 0 reveals an absence of double COs and is thus the presence of CO interference. The crossover interference is undetectable in both female and male meiosis of hyper recombination mutants with ZYP1 and/or RECQ4 mutations, consistent with previous studies (Séguéla-Arnaud et al. 2015; Serra et al. 2018; Fernandes et al. 2018; France et al. 2021; Capilla-Pérez et al. 2021; Durand et al. 2022).

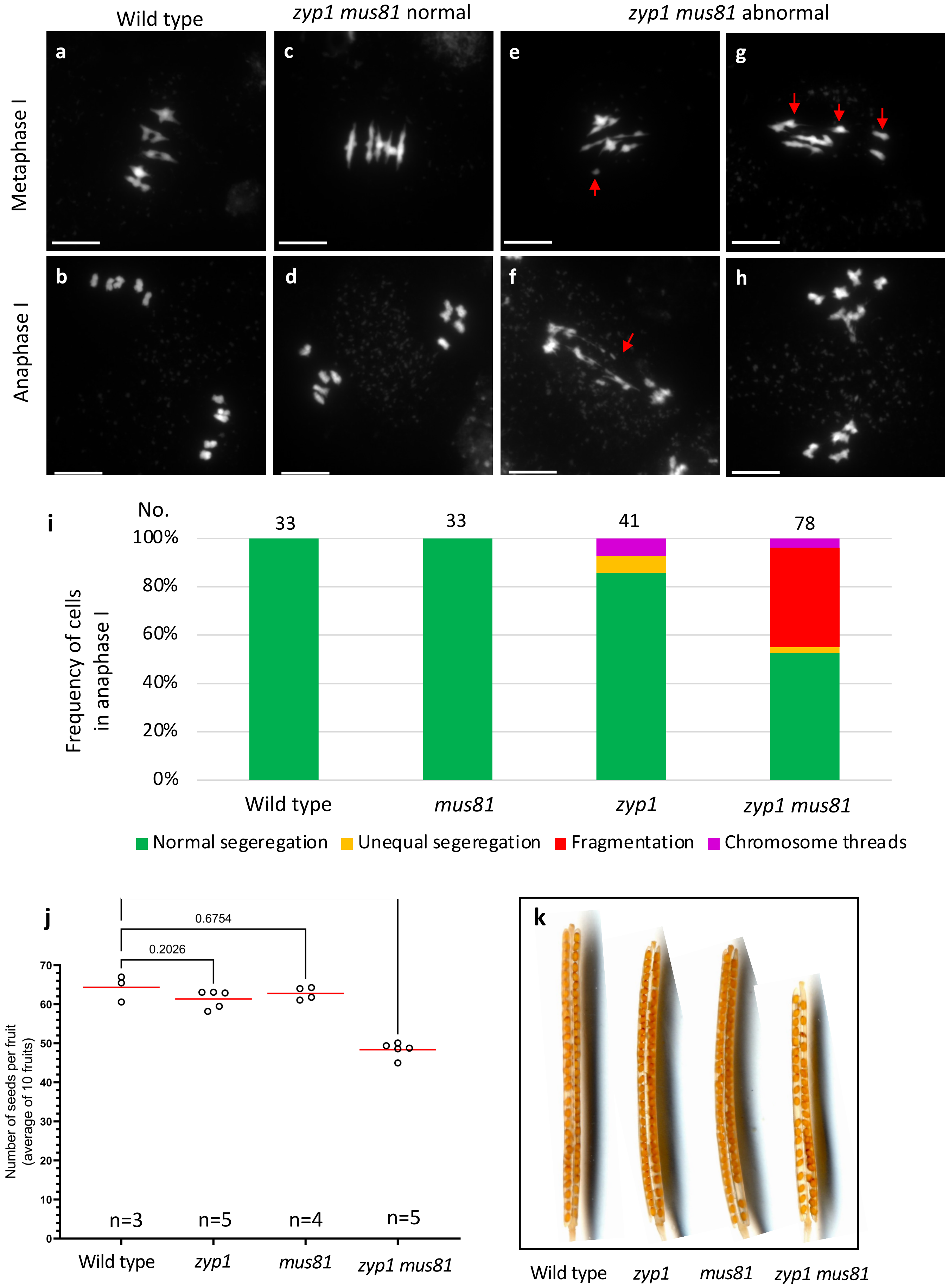

Wild type
zyp1 mus81 normal
zyp1 mus81 abnormal
Metaphase I
Anaphase I
a
b
c
d
e
f
g
h
i
No.
33
33
41
78
Frequency of cells in anaphase I
100%
80%
60%
40%
20%
0%
Wild type
mus81
zyp1
zyp1 mus81
Normal segeregation
Unequal segeregation
Fragmentation
Chromosome threads
j
0.6754
0.2026
Number of seeds per fruit (average of 10 fruits)
70
60
50
40
30
20
10
0
n=3
n=5
n=4
n=5
Wild type
zyp1
mus81
zyp1 mus81
k
Wild type
zyp1
mus81
zyp1 mus81

**Fig S6. Analysis of meiosis and fertility in wild type, *zyp1*, *mus81* and *zyp1 mus81*.**

**a-g**. DAPI-stained meiotic chromosome spreads from male meiocytes in wild type (**a, b**), *zyp1*(**c, d**), *mus81* (**e,f**) and *zyp1 mus81* (**g, h**). **a, c, e, g** Metaphase I, **b, d, f, h** Anaphase I. Red arrows pointed out abnormal chromosome connections, fragments and chromosome threads. **i.** Quantification of different chromosome behaviors at metaphase I in wild type, *zyp1*, *mus81* and *zyp1 mus81* in Col background. Cells were categorized according to normal (5 bivalents) and abnormal chromosome behavior (unequal segregation, fragmentation and chromosome threads). The number of analyzed cells are indicated above the bar. **j.** Quantification of fertility. Each dot represents the fertility of an individual plant, measured as the number of seeds per fruits averaged on ten fruits. The red bar shows the mean. All plants were grown in parallel, and the wild-type controls are siblings of the mutants. The number n of analyzed plants is indicated and p values are one-way ANOVA followed by Fisher's LSD test. **k.** Representative cleared fruits of wild type, *zyp1*, *mus81* and *zyp1 mus81* in Col background.

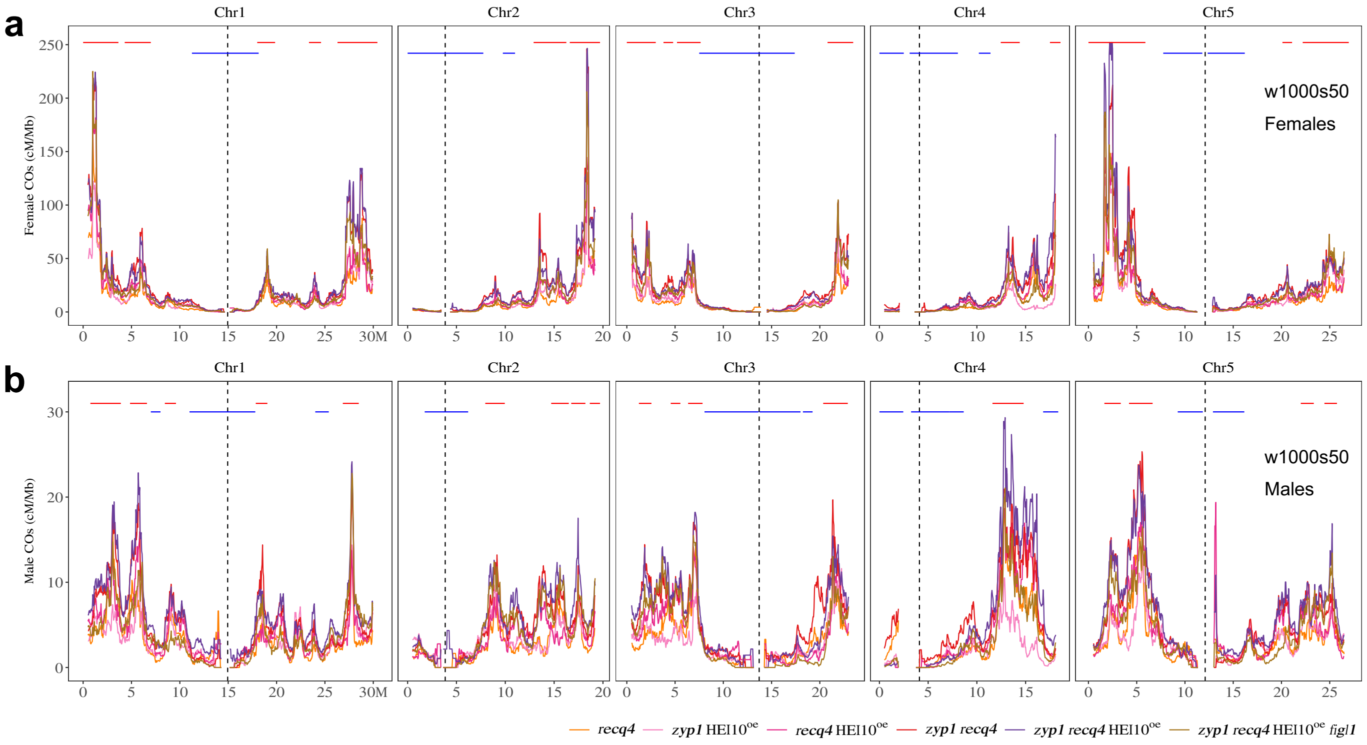


**Fig S7. The fold-change analysis of CO frequencies along chromosomes in females and males.**

The chromosomal distribution of fold-changes of CO frequencies (sliding window-based, window size 1 Mb, step size 50 kb) against wild-type in females (**a.**) and males (**b.**), respectively. The hot and cold zones of CO burst from hyper recombination mutants were colored by red and blue horizontal lines, separately.

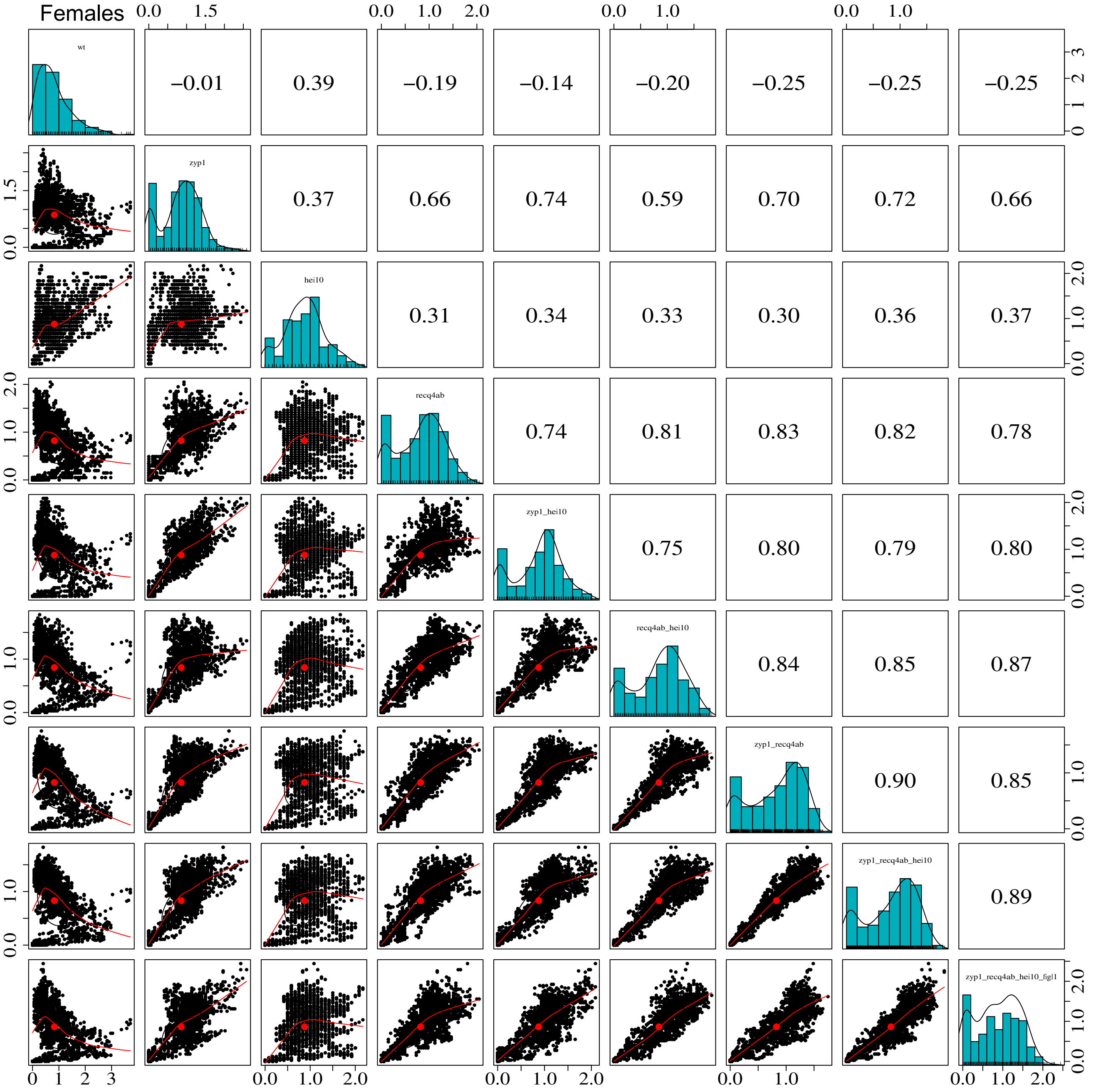


**Fig S8. Correlation analysis of the relative distribution of COs in females.**

Spearman's correlation between each pair of genotypes is shown in the upper triangle panel. The chromosome 4 is ignored in the analysis.

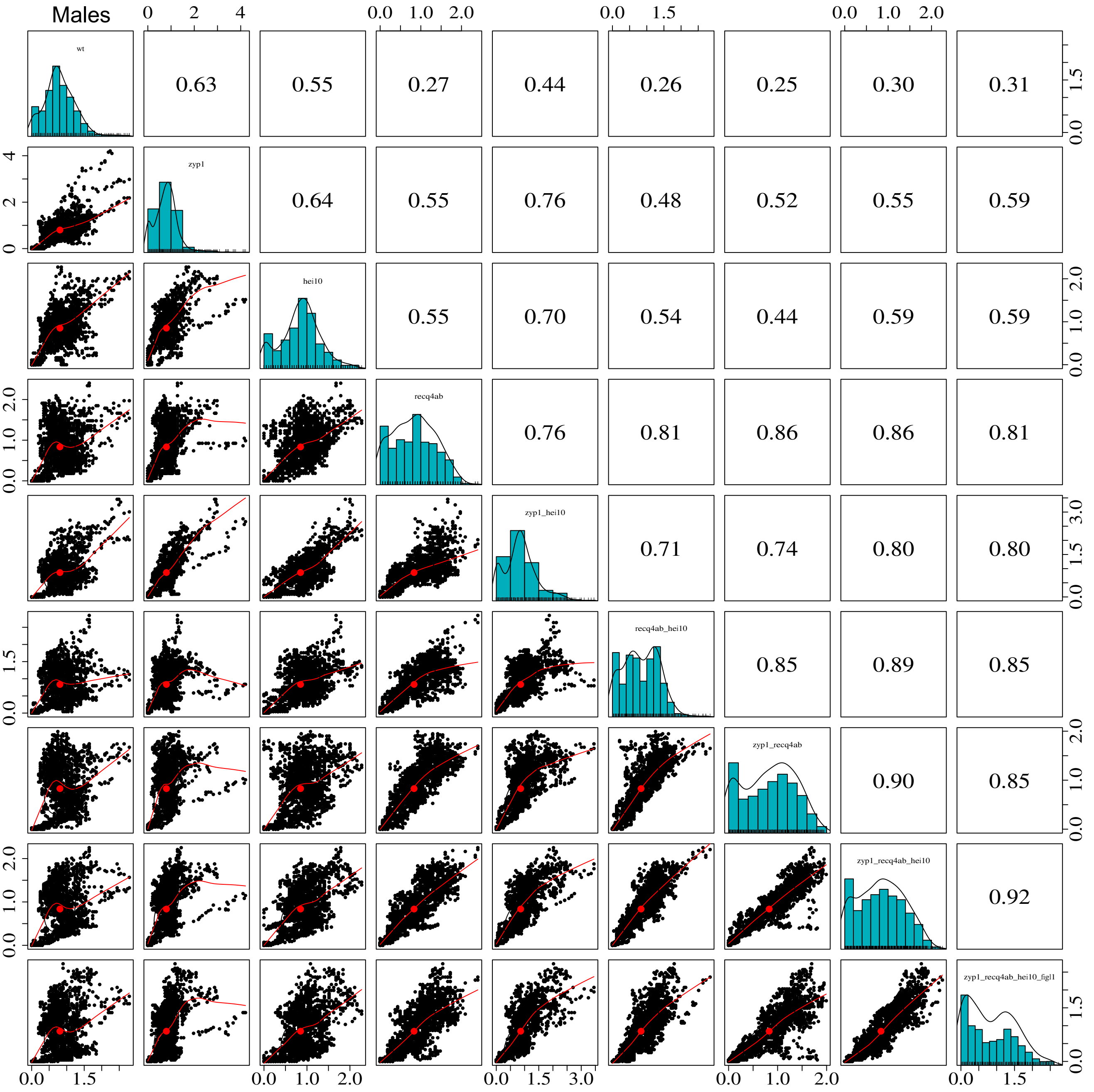


**Fig S9. Correlation analysis of the relative distribution of COs in males.**

Spearman's correlation between each pair of genotypes is shown in the upper triangle panel. The chromosome 4 is ignored in the analysis.

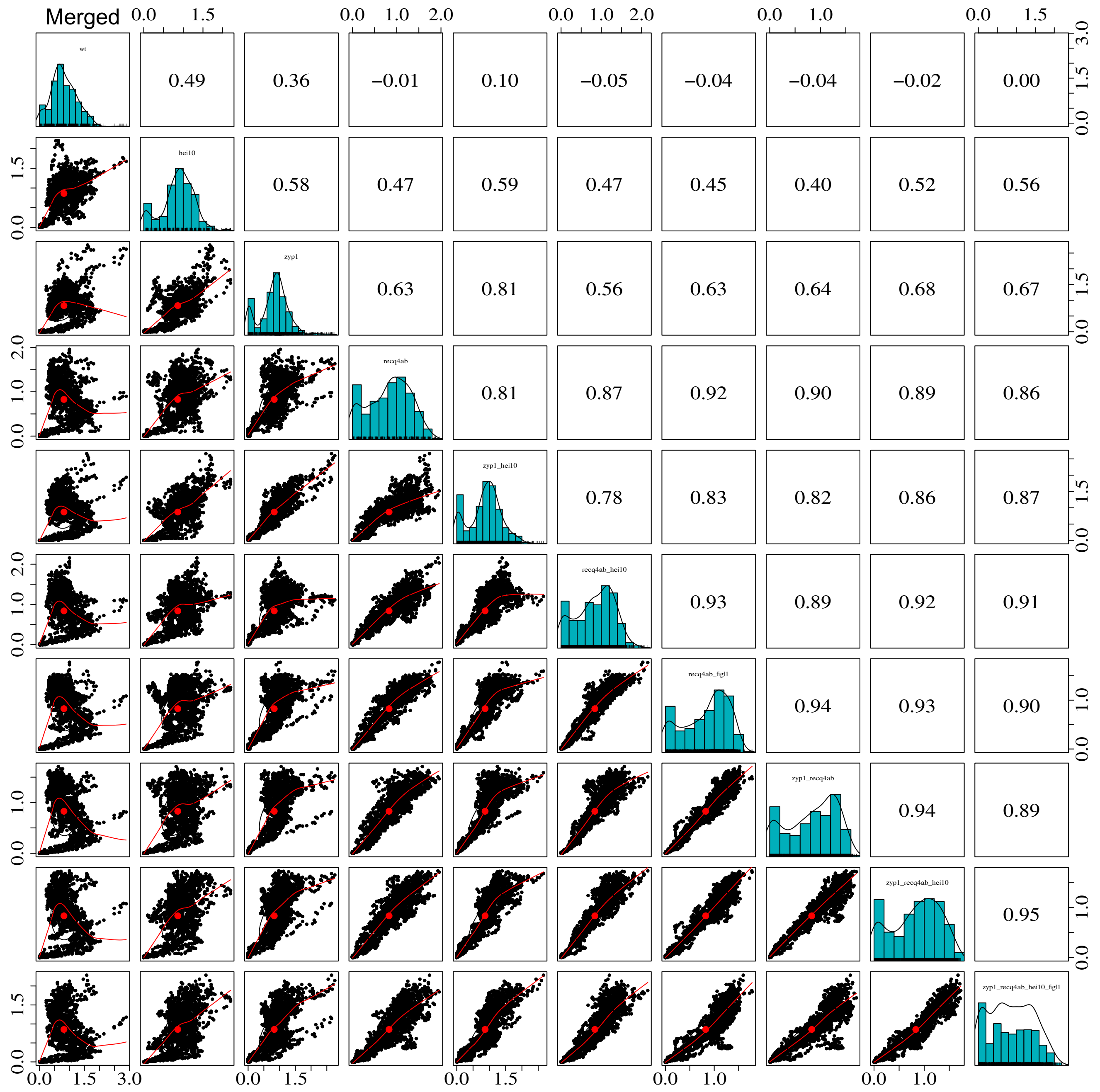


**Fig S10. Correlation analysis of the relative distribution of COs in F2s or pseudo F2.**

Spearman's correlation between each pair of genotypes is shown in the upper triangle panel. The chromosome 4 is ignored in the analysis.

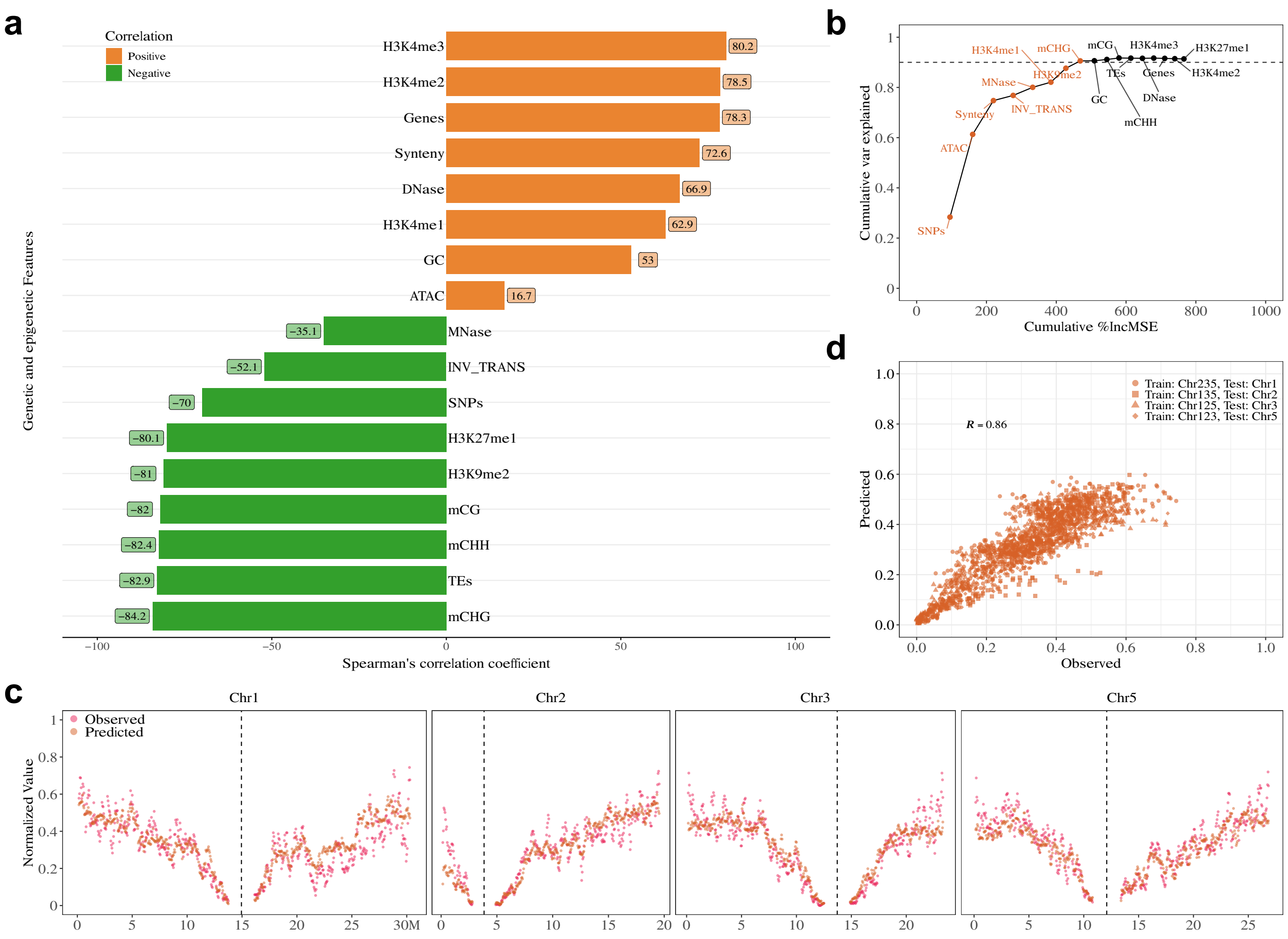


**Fig S11. Association and prediction of precursor distribution with genetic and epigenetic features on chromosome arms.**

**a.** Spearman's correlation test shows the comparison with features along chromosome arms, with differences in colour and length according to the correlation scale. SNPs (SNPs density between Col and L*er*), INV_TRANS (inversions and translocations between Col and L*er*), Synteny (collinearity between Col and L*er*), Genes, TEs and GC (expressed gene in meiocytes, TE and GC density), ATAC and DNase (chromatin accessibility, ATAC-seq and DNase-seq, log2(Tn5/gDNA) and log2(DNase/gDNA)), H3K4me1/2/3, H3K9me2, H3K27me1 (euchromatin, heterochromatin, and Polycomb histone marks, ChIP-seq, log2(ChIP/input)), mCG, mCHG and mCHH (DNA methylation in CG, CHG, and CHH contexts, proportion methylated cytosine), MNase (nucleosome occupancy, MNase-seq, log2(MNase/gDNA)). **b.** The cumulated proportion of variation that can be explained with the features at the chromosome arm scale. The top eight most important features are coloured, for which the cumulative proportion of variation that can be explained reaches the plateau. **c.** The chromosomal distribution of observed and predicted precursor maps. The precursor profiles of individual chromosomes were predicted using profiles of the top eight most important features from the other three chromosomes. **d.** The Spearman's correlation test between the predicted and observed precursor distributions. The training-testing dataset is differentiated in shapes.
